# Cell-type specific *in vitro* gene expression profiling of stem-cell derived neural models

**DOI:** 10.1101/2020.04.30.064709

**Authors:** James A. Gregory, Emily Hoelzli, Rawan Abdelaal, Catherine Braine, Miguel Cuevas, Madeline Halpern, Natalie Barretto, Nadine Schrode, Güney Akbalik, Kristy Kang, Esther Cheng, Kathryn Bowles, Steven Lotz, Susan Goderie, Celeste M. Karch, Sally Temple, Alison Goate, Kristen J. Brennand, Hemali Phatnani

## Abstract

Genetic and genomic studies of brain disease increasingly demonstrate disease-associated interactions between the cell types of the brain. Increasingly complex and more physiologically relevant human induced pluripotent stem cell (hiPSC)-based models better explore the molecular mechanisms underlying disease, but also challenge our ability to resolve cell-type specific perturbations. Here we report an extension of the RiboTag system, first developed to achieve cell-type restricted expression of epitope-tagged ribosomal protein (RPL22) in mouse tissue, to a variety of *in vitro* applications, including immortalized cell lines, primary mouse astrocytes, and hiPSC-derived neurons. RiboTag expression enables efficient depletion of off-target RNA in mixed species primary co-cultures and in hiPSC-derived neural progenitor cells, motor neurons, and GABAergic neurons. Nonetheless, depletion efficiency varies across independent experimental replicates. The challenges and potential of implementing RiboTags in complex *in vitro* cultures are discussed.

## Introduction

The many cell types of the brain interact to influence each other’s function in health and disease^1^. Moreover, the growing list of rare and common risk loci associated with brain disease implicate a variety of cell types across disorders, with genetic risk sometimes enriched in cell types previously not thought to be related to disease pathology^2^. Non-neuron cell-autonomous effects are increasingly recognized to play a critical role across psychiatric and neurodegenerative disease^3^. Human induced pluripotent stem cell (hiPSC)-based models represent a powerful approach to study the complex etiology of brain disease^4^. Towards this, it is necessary to assemble increasingly complex *in vitro* models that better capture neuronal circuitry^5^, astrocyte support^6^, myelination^7^, microglia^8^, vasculature^9^, and blood-brain-barrier functions^10^. However, interrogating cell-type-specific processes within increasingly complex networks, particularly of cell types represented at low abundance, is difficult using conventional approaches.

Complementary physical, biochemical, and computational approaches have been developed to identify cell-type specific transcriptomes. Physical methods for isolating cell-type specific RNA include fluorescence activated cell sorting (FACS)-based enrichment^11,12^, single cell RNA sequencing (scRNA-seq)^13,14^, and laser capture microdissection (LCM)^15–17^. Single-cell suspensions required for FACS and scRNA-seq can be challenging to prepare for highly arborized neural cells. scRNAseq provides cellular resolution data at low read depth and despite recent advances^18,19^, remains prohibitively expensive for large scale studies. Low read depth and other technical considerations can bias scRNA-seq studies, with certain cell types or transcripts over-represented or missing in the sequencing libraries generated through these methods^20^. LCM is labor intensive, requires highly specialized equipment, and has similar limitations with respect to transcript diversity^21^. Computational approaches to infer cell type abundance and deconvolve cell-type specific transcriptomes from bulk RNA-seq^22^ are improving as scRNA-seq reference datasets become more widely available across cell types^23^.

RiboTag^24^ and translating ribosome affinity purification (TRAP)^25,26^ are similar biochemical approaches that rely on cell-type specific expression of epitope-tagged ribosomal subunits to biochemically enrich mRNA from cell types of interest. RiboTag and TRAP rely on the tight interaction between the ribosome and mRNA to indirectly immunoprecipitate translated RNA via RPL22 or RPL10a, respectively. They do not require specialized equipment, dissociation to single cells, and these methods have been used extensively for studies of murine neurons^27,28^, microglia^29^, astrocytes^30^, and macrophages^31^. We reasoned that the same strategy could be used to purify cell-type specific mRNA from heterogeneous *in vitro* co-cultures. Additionally, mRNA from multiple cell types could be simultaneously isolated if each cell type expressed a ribosomal subunit with a different epitope tag.

Here we tested RPL22 fused V5^32^, hemagglutinin^33^, or Flag^34^ affinity-tags for immunoprecipitation of cell type specific mRNA *in vitro;* unexpectedly, only RPL22-HA and RPL22-V5 RiboTags co-immunoprecipitated mRNA. We demonstrate that RPL22-HA and RPL22-V5 immunoprecipitation efficiently enriched mRNA from mixed species co-cultures of immortalized cell lines. When expressed in hiPSC-derived motor neurons (MNs), neural progenitor cells (NPCs), and induced GABAergic neurons (iGANs), immunoprecipitated RNA was enriched for protein coding genes with a concomitant decrease in lncRNA and mitochondrially encoded genes. Unfortunately, across independent experimental replicates, cell-type specific mRNA enrichment was highly variable in heterocellular cultures composed of hiPSC-derived neurons and/or primary mouse astrocytes.

## Results

### Construction of epitope tagged RPL22

C-terminal HA-tagged RPL22 has been extensively used to purify cell-type specific RNA from mouse CNS tissue^24^. We reasoned that RPL22 may tolerate additional epitope tags and, when used in combination, could be used to biochemically separate RNA from multiple cell types in a mixed co-culture if each cell type expresses a uniquely tagged RPL22. We fused *RPL22* to three well-established epitope tags (V5, Flag, HA) and cloned them into multicistronic lentiviral vectors with antibiotic resistance genes and/or fluorescent proteins (FPs) separated by picornavirus 2A peptides (Fig. S1). All three C-terminal RPL22 fusions (hereafter referred to as RiboTags) accumulate in HEK293 cells when transcription is controlled by the EF1-a promoter (Fig. 1A-C – top panels). RPL22-1xV5 (hereafter RPL22-V5), RPL22-1xFlag, and RPL22-3xHA (hereafter RPL22-HA), migrate slightly above the predicted molecular weights of 16.6, 16.1, and 20.5 kDa, respectively. RNA was consistently recovered following immunoprecipitation (IP) of RPL22-HA or RPL22-V5. Conversely, RNA yields following RPL22-1xFlag or RPL22-3xFlag IP were inconsistent, resulting in minimal to no mRNA recovery. Lack of mRNA recovery, despite RPL22-Flag protein accumulation, suggests that the Flag and 3xFlag epitopes may interfere with incorporation of RPL22 into the ribosome. Consistent with this hypothesis, RPLP0 co-immunoprecipitated with RPL22-HA and RPL22-V5, but not RPL22-Flag (Fig 1A-C – bottom panels) or RPL22-3xFlag (data not shown).

**Figure 1.**
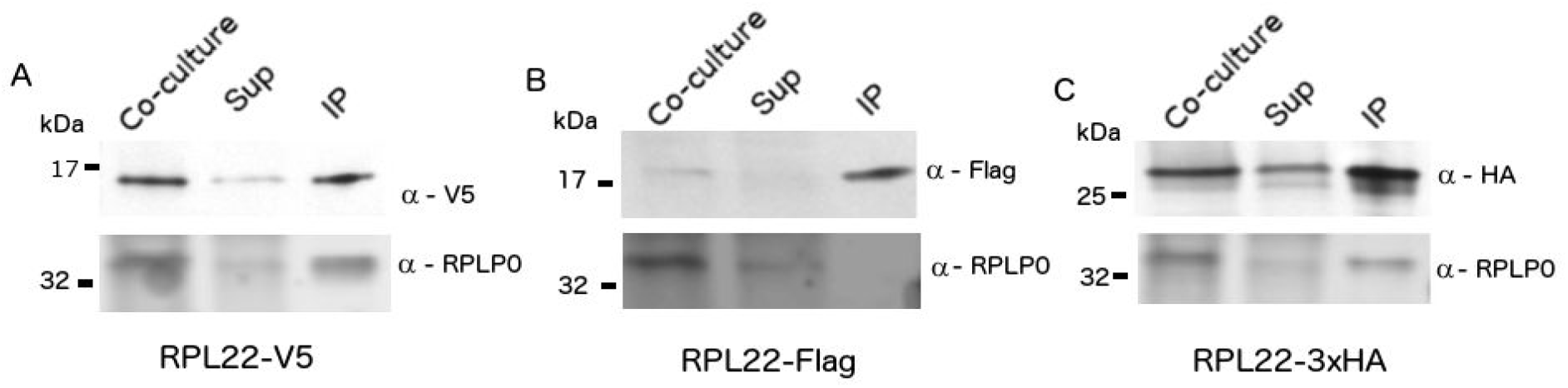
RiboTag immunoblot of HEK293 cells expressing RiboTag constructs. HEK293 cells were transduced with lentivirus encoding (A) RPL22-V5, (B) RPL22-Flag, or (C) RPL22-3xHA. Total soluble protein from co-culture lysates, supernatant (Sup), and immunoprecipitations (IP) were resolved by SDS-PAGE, transferred to nitrocellulose, and detected with antibodies to RPLP0 and V5, Flag, or haemagglutinin (HA).

### Biochemical separation of cell-type specific RNA in co-cultures using RiboTags

To tightly control RiboTag expression, we constructed lentiviral vectors encoding mCherry-T2A-RPL22-V5 and GFP-T2A-RPL22-HA driven by the tetracycline response element (P_TRE_). Additionally, we moved RPL22 to the 3’ end of the multicistronic construct to eliminate extra amino acids in the epitope tag resulting from cleavage of the 2A peptide (Fig. S2). When used in combination with reverse tetracycline-controlled transactivator (rtTa), RiboTag expression is induced with doxycycline. We generated stable RiboTag HEK293 cell lines via lentiviral transduction and doxycycline induction, followed by FACS. The RPL22-HA and RPL22-V5 expressing cell lines were seeded together in approximately equal numbers. The mixed co-culture was split in half twenty-four hours post-doxycycline induction; half was used for HA IP and the other for V5 IP. GFP and mCherry accumulation were confirmed by microscopy and Western blot (Fig. S3A-B). RPL22-HA and RPL22-V5 protein accumulation were confirmed by Western blot using anti-HA and anti-V5 antibodies, respectively (Fig. 2A). RPL22-HA appears as a doublet following IP, suggesting possible degradation. Absolute *GFP* and *mCherry* mRNA abundance was measured by reverse transcription digital PCR (RT-dPCR; Table S3, Fig. 2B). Each cell expresses *GFP* or *mCherry,* allowing relative levels to be used as a proxy for cell-type specific mRNA abundance. The initial *mCherry/GFP* ratio in mRNA isolated from the co-culture was approximately 9:2 (Fig. 2B – yellow). As expected, this ratio is increased in mRNA isolated with V5 antibodies and decreased in mRNA isolated with HA antibodies (Table S4; Fig. 2B). The combined effect was an 88-fold difference in the *mCherry/GFP* ratio between the HA and V5 IP RNA samples.

**Figure 2.**
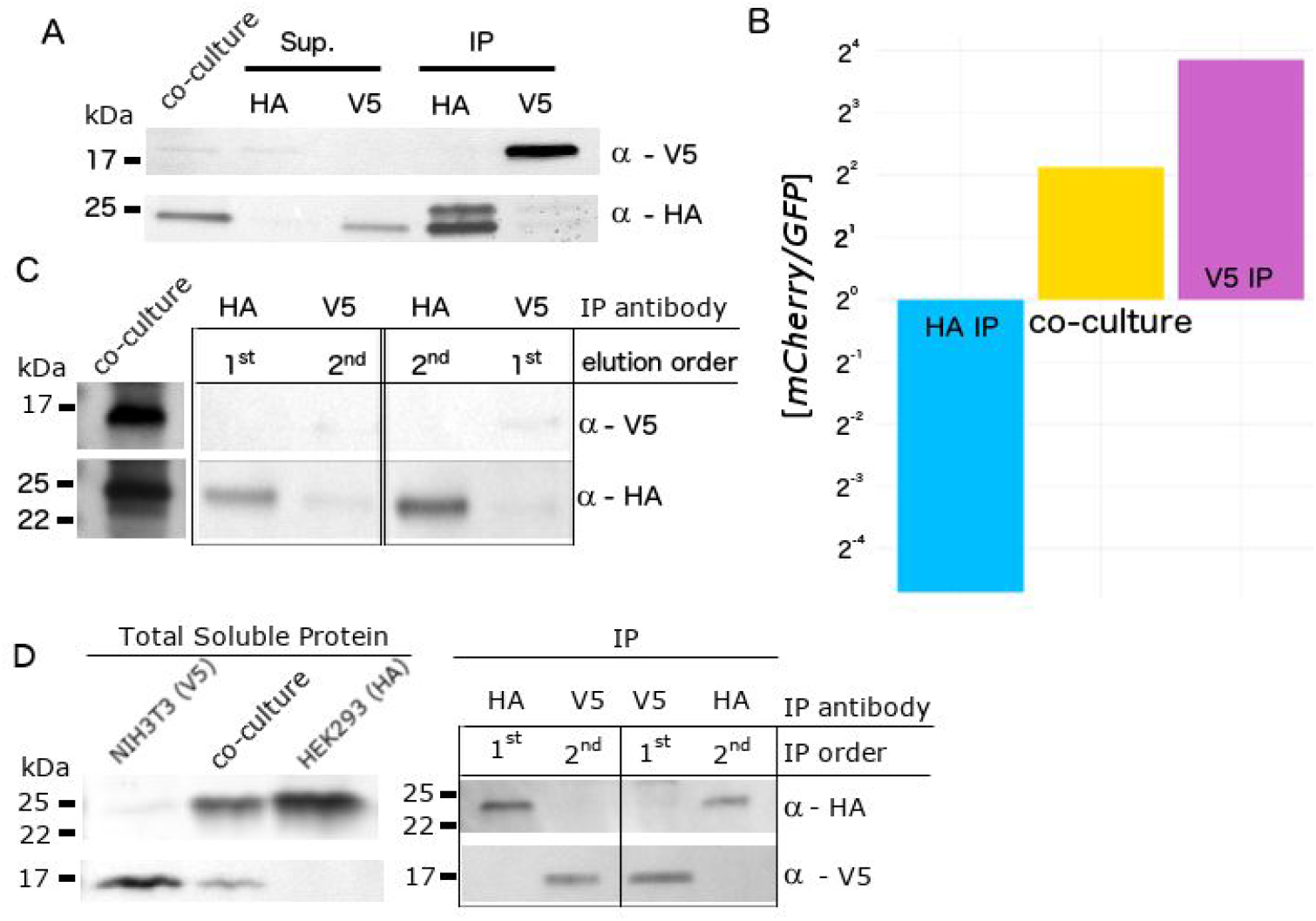
Co-culture of HEK293 cells expressing RPL22-HA or RPL22-V5. (A) Total soluble protein from co-culture lysates, supernatants (Sup), and immunoprecipitated (IP) protein were resolved by SDS-PAGE, transferred to nitrocellulose, and probed with antibodies to HA or V5. (B) mCherry and GFP mRNA concentration was measured in triplicate by reverse transcription digital PCR in RNA isolated from the co-culture lysates (yellow), HA IP (blue), and V5 IP (purple), and plotted as a ratio. (C) RPL22-V5 and RPL22-HA were simultaneously IPed from co-culture lysates and selectively eluted in the indicated order using V5 or HA peptides, respectively. Total soluble protein from co-culture lysate and IPed protein were resolved by SDS-PAGE, transferred to nitrocellulose, and probed with antibodies to V5 or HA. (D) RPL22-V5 and RPL22-HA were sequentially IPed in the indicated order from co-culture lysates. Total soluble protein from co-culture lysates and IPed protein were resolved by SDS-PAGE, transferred to nitrocellulose, and probed with antibodies to HA or V5.

In the above experiments, co-culture lysates were split equally for HA IP and V5 IP. This strategy reduces the potential RNA yield by half, which is problematic for precious samples or when obtaining large numbers of cells is impractical. We reasoned that immunoprecipitation with HA and V5 antibodies from the same sample could improve RNA recovery. To test this hypothesis, we implemented two strategies. First, we simultaneously immunoprecipitated RPL22-HA and RPL22-V5 using antibody-conjugated magnetic beads followed by sequential elution with HA then V5 peptides or vice versa. We observed incomplete elution of RPL22-V5 using V5 peptides and RPL22-V5 contamination in the HA peptide eluted fraction (Fig. 2C). Second, we tested a sequential IP strategy whereby magnetic beads conjugated to only one antibody is used for IP and the resulting supernatant is used for a second IP with the other antibody. Each IP fraction was tested for the presence of RPL22-HA or RPL22-V5 by Western blot (Fig. 2D). RPL22-HA and RPL22-V5 were only detected in IP fractions using HA or V5 antibodies, respectively, regardless of the order that the IPs were performed. Thus, sequential IP can be used to separate epitope-tagged RPL22 from distinct cell populations.

### Transcriptome-wide cell-type specific enrichment in mixed species co-cultures using RiboTags

To assess cell-type specific enrichment transcriptome-wide, we co-cultured human (HEK293) and mouse (NIH-3T3) cells transduced with P_TRE_-GFP-T2A-RPL22-HA and P_TRE_-mCherry-T2A-RPL22-V5, respectively. IPs were carried out sequentially using both possible orders (e.g. HA then V5 and vice versa). We observed a greater than 1000-fold difference in the *mCherry/GFP* ratio, as measured by RT-dPCR, between the HA and V5 IP mRNA regardless of the order that the IPs were performed (Fig. 3A, Table S5). For both HA and V5 IPs, the RNA yield was reduced in the second IP (Table S6). We next used RNA-seq to evaluate transcriptome-wide cell-type specific mRNA enrichment (Fig. 3B). Reads were assigned as human or mouse using STAR^35^ by aligning all reads to a combined reference genome consisting of the canonical chromosomes from GRCh38 (human) and GRCm38 (mouse) and sequences for *GFP* and *mCherry. GFP* and *mCherry* read counts were consistent with RT-dPCR results (Fig. S4, Table S7). The initial species composition of reads in the mixed co-culture was 72% mouse and 28% human (Fig. 3B; Table S7). IP using HA or V5 antibodies increased the read composition of human and mouse reads, respectively, and the species composition was similar regardless of the order that the IPs were performed. As expected, gene expression from HEK293 monocultures correlates with HA IP samples whereas V5 IP samples more closely correlate with NIH 3T3 monocultures (Fig. 3C).

**Figure 3.**
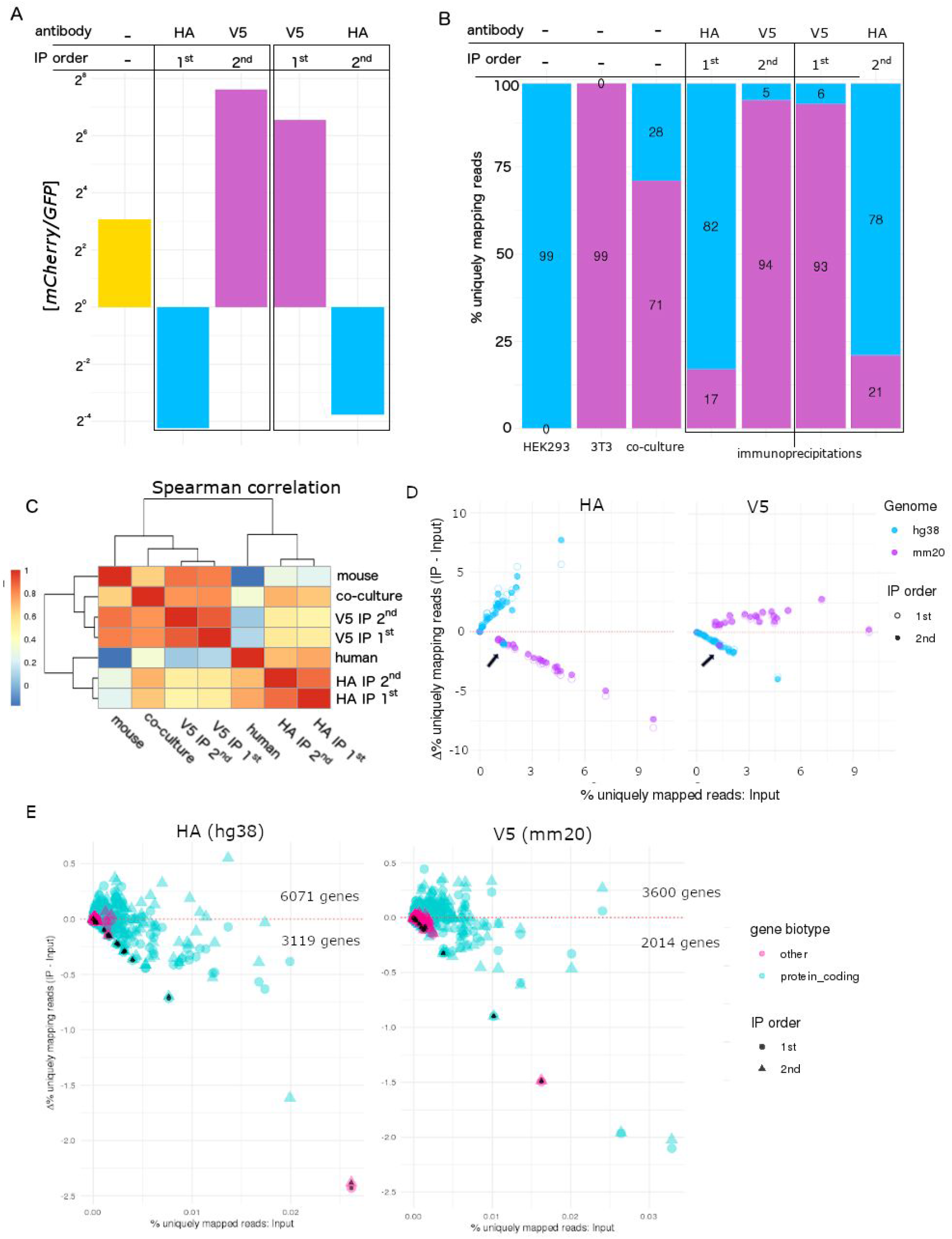
Application of RiboTags to mixed species co-cultures of HEK293 cells expressing RPL22-HA and NIH3T3 cells expressing RPL22-V5. (A) mCherry and GFP mRNA concentration was measured in triplicate by reverse transcription digital PCR in RNA isolated from co-culture lysates (yellow), HA IP (blue), and V5 IP (purple), and plotted as a ratio. (B) RNA-seq reads were mapped to a hybrid reference genome containing hg38 and mm20 chromosomes and quantified by species for monocultures, co-culture, and IP samples. (C) Spearman correlation plot of RNA-seq for co-culture and IP samples. (D) The change in uniquely mapping reads (IP – input) for each chromosome was plotted against the initial (Input) percent uniquely mapping reads. Each dot represents a chromosome (hg38 – blue, mm20 – purple). First and second IPs are represented by open and filled dots, respectively. Chromosomes that fall above the red dotted line are enriched whereas chromosomes that fall below the red dotted line were depleted in IP samples compared to the Input. (E) The change in on-target (e.g. hg38 for HA IP and mm20 for V5) percent uniquely mapping reads (IP – Input) was plotted against the initial (Input) percent uniquely mapping reads at the gene level. Protein coding genes are in cyan and non-protein coding genes in pink. First and second IPs are represented by circles and triangles, respectively. Mitochondrially encoded genes have additional black colored fill.

There was considerable cross species contamination despite species specific expression of HA or V5-tagged RPL22. For RNA isolated by HA IP, we considered reads that uniquely mapped to the human genome as on-target and uniquely mapped mouse reads as off-target; we inverted the analysis for V5 IPs. We quantified the number of reads mapping to each chromosome (hg38 and mm20) before and after IP and plotted the change (e.g. input – IP) for each chromosome (Fig. 3D). Except for the mitochondrial genome, (Fig. 3D – arrows), which are not translated by RPL22-containing ribosomes, chromosomes with on-target reads were enriched and those with off-target reads were depleted. When RPL22-HA is expressed in HEK293 cells, HA IP resulted in enrichment of reads mapping to hg38 (Fig. 3D – HA; blue dots above the x-axis). Similarly, when RPL22-V5 expression is restricted to NIH3T3 cells, V5 IP enriched for reads mapping to mm20 chromosomes (Fig. 3D – V5; purple dots above the x-axis). We observed a striking negative correlation for the depletion of off-target reads (R^2^ – 0.99; Table 1), the negative slope of which we define as the depletion efficiency (0.74 – 0.85; Table 1). The maximum theoretical depletion efficiency is 1 (see materials and methods). We performed a similar analysis for on-target reads at the gene level (Fig. 3E). As expected, mitochondrially encoded genes were depleted in a manner similar to off-target genes in both cases (Fig. 3E – black fill). The relative change (IP – input) varied considerably across on-target protein coding genes. More protein coding genes were enriched than depleted. Conversely, non-protein coding genes were primarily depleted.

**Table 1.** Relative enrichment of species specific mRNA in mouse/human co-cultures using RiboTags. The change in uniquely mapping reads (IP – input) for each mouse and human chromosome was plotted against the initial (Input) percent uniquely mapping reads Below is the species-specific linear regression of the change in uniquely mapping reads from (% unique reads vs Δ unique reads) on a chromosome level. ‘Species’ indicates which chromosomes were used to calculate the regression and ‘off/on target’ indicates whether RPL22 was expressed in that species (on-target) or not (off-target).

| Sample | IP Order | On target species | Off target species | slope | R<sup>2</sup> | p-value | RNA yield (ng) |
| --- | --- | --- | --- | --- | --- | --- | --- |
| V5-1 (3T3) | 1 | human | off target | -0.8 | 0.99 | p < 0.001 | 388 |
|  | 1 | mouse | on target | 0.12 | 0.08 | p < .10 |  |
| V5-2 (3T3) | 2 | human | off target | -0.85 | 0.99 | p < 0.001 | 102 |
|  | 2 | mouse | on target | 0.14 | 0.12 | p < 0.06 |  |
| HA-1 (HEK293) | 1 | human | on target | 1.26 | 0.59 | p < 0.001 | 200 |
|  | 1 | mouse | off target | -0.81 | 0.99 | p < 0.001 |  |
| HA-2 (HEK293) | 2 | human | on target | 1.56 | 0.74 | p < 0.001 | 67 |
|  | 2 | mouse | off target | -0.74 | 0.99 | p < 0.001 |  |

### Application of RiboTag to hiPSC-derived neural cell types

The expanding repertoire of directed differentiation and neural induction protocols allows *in vitro* studies of a variety of human neural cells from donor specific genetic backgrounds. We constructed stable hiPSC and neural progenitor cell (NPC) lines using lentiviral vectors encoding RiboTags. GFP or mCherry accumulation diminished over long-term culture in NPCs that constitutively express epitope-tagged RPL22. We also observed low and inconsistent P_TRE_-RiboTag expression in MNs derived from hiPSCs that were transduced and purified by flow cytometry. Thus, long-term culture and differentiations may be associated with gene silencing. Lentiviral transduction of P_TRE_-RiboTags during neural induction or in differentiated neurons improved RiboTag expression. Marker gene expression was confirmed by RNA-seq (Fig. 4A) for hiPSC-derived MNs (n=7 input; n=3 IP), GABAergic neurons (iGANs; n=2 input; n=2 IP), and NPCs (n=1 input; n=1 IP). We confirmed RPL22 immunoprecipitation by Western blot (Fig. 4B). Using Ensembl gene biotypes to categorize annotated genes, we observed a significant enrichment of protein coding transcripts and concomitant decrease in long non-coding RNA (lncRNA) and pseudogenes (Fig. 4C). In contrast, mitochondrial protein coding transcripts were depleted in IP samples compared to input controls (Fig. 4D). Thus, RiboTags can be expressed in hiPSC-derived cells and used to enrich for translated mRNA by IP.

**Figure 4.**
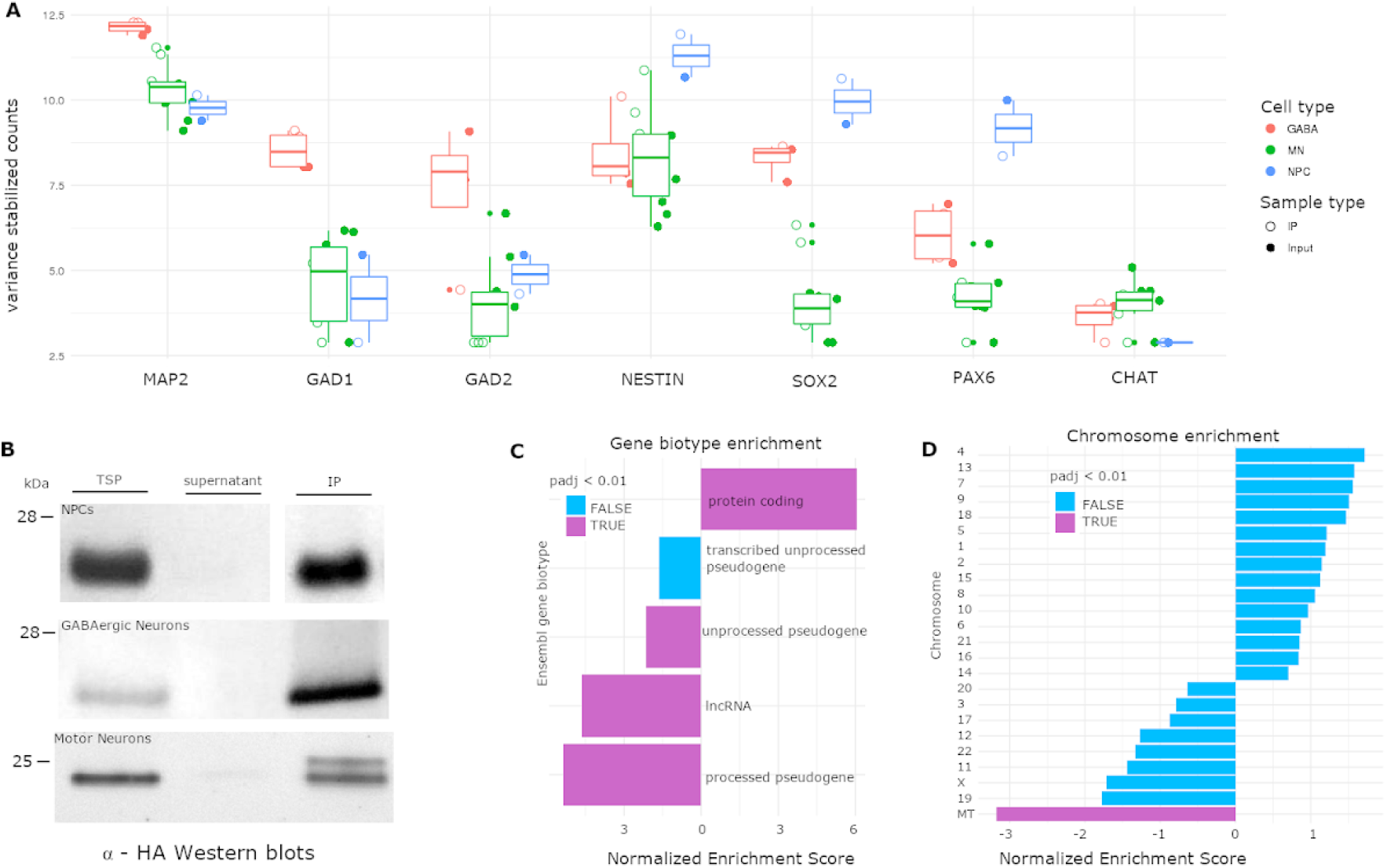
Application of RiboTags to hiPSC-derived neural progenitor cells (NPCs), GABAergic neurons (iGABAs), and motor neurons (MNs). (A) Variance stabilized counts of neural (MAP2), GABAergic (GAD1, GAD2), neural progenitor (NESTIN, SOX2, PAX6), and cholinergic (CHAT) marker genes from hiPSC-derived NPCs (blue), iGABAs (salmon), and MNs (green) from lysates (filled dots) and IP (open dots) samples using RNA-seq. (B) Total soluble protein (TSP) from cell lysates, supernatants (Sup), and immunoprecipitated (IP) protein were resolved by SDS-PAGE, transferred to nitrocellulose, and probed with antibodies to HA for hiPSC-derived NPCs, iGABAs, and MNs. (C) Gene set enrichment analysis (GSEA) using Ensembl gene biotypes across all annotated genes. (D) GSEA of protein coding genes grouped by chromosomes. TSP – total soluble protein. IP – immunoprecipitate.

We next applied P_TRE_-RiboTag to co-cultures of hiPSC-derived MNs and primary mouse astrocytes. Cells were transduced with P_TRE_-RiboTag (HA or V5) lentivirus as monocultures, mixed, and co-cultured for two weeks. We performed six independent replicates (e.g. independent motor neuron differentiations, mouse astrocyte dissections, and transductions) using mixed species cocultures to evaluate the variability between experiments. RNA from co-culture (n=6), MN IPs (n=6), and mAstro IPs (n=3) was isolated 24-hrs post-doxycycline treatment for RNA-seq. Reporter gene expression was not significantly different between RiboTags or cell type (Fig. S5A-B). Initial co-culture species composition varied by as much as 50% (Fig. 5A, Fig. S5C) and enrichment of GFP or mCherry relative to co-cultures ranged from 2.4 to 194-fold (Fig. S5D). Similar to co-cultures using HEK293 and NIH3T3 cells, we observed an enrichment of reads mapping to on-target chromosomes and depletion of reads mapping to off-target chromosomes compared to the co-culture (Fig. 5A; Fig. S6). Depletion efficiency ranged from 0.25-0.84 and 0.46-0.87 for RPL22-V5 and RPL22-HA, respectively (Fig. 5B). The correlation between the percentage of reads mapping to off-target chromosomes in pre-vs post-IP samples was reduced for low efficiency IPs (Fig. 5B, Fig. S6. Table 2). Reporter gene expression (GFP/mCherry), which is a proxy for RiboTag expression, and initial co-culture composition did not correlate with depletion efficiency (Fig. S7). Thus, differential RiboTag expression in mixed co-cultures allows for cell-type specific RNA enrichment, but the enrichment is not consistent across replicates.

**Figure 5.**
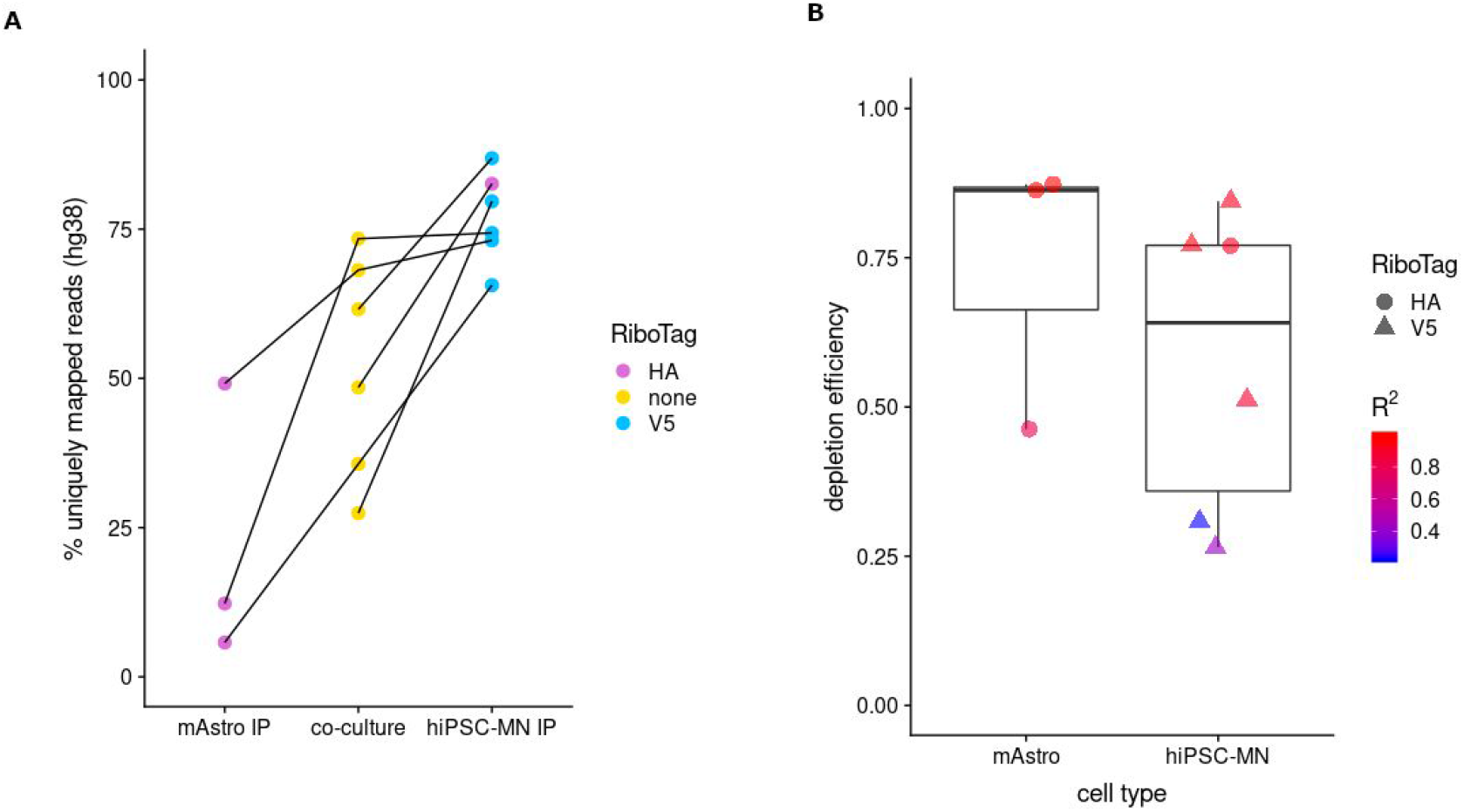
Application of RiboTags to co-cultures of hiPSC derived motor neurons and primary mouse astrocytes. (A) RNA-seq reads were mapped to a hybrid reference genome containing hg38 and mm20 chromosomes and protein coding genes were quantified by species for co-cultures (yellow), HA IP (purple), and V5 IP (blue). Lines indicate matched samples (e.g. IPs came from the indicated co-culture). (B) The depletion efficiency was calculated for HA (circles) and V5 (triangles) RiboTag samples using off-target reads (mm20 for hiPSC-MN IPs; hg38 for mAstro IPs). Color indicates the correlation between the IP sample and the matched co-culture.

**Table 2.** Relative depletion of off-target mRNA in co-cultures of hiPSC-motor neurons and primary mouse astrocytes. The change in uniquely mapping reads (IP – input) for each mouse and human chromosome was plotted against the initial (Input) percent uniquely mapping reads. Below is the species-specific linear regression of the change in uniquely mapping reads from (% unique reads vs Δ unique reads) on a chromosome level for the off-target species (e.g. only relative depletion is shown).

| Library | Input_library | MN_differentiation | IP_Cell_Type | RiboTag | species | slope | R_squared | p_value | RNA yield (ng) |
| --- | --- | --- | --- | --- | --- | --- | --- | --- | --- |
| 5 | 2 | 2019-06-03 | mAstro | HA | mouse | -0.46 | 0.82 | 0.001 | 593.6 |
| 8 | 2 | 2019-06-03 | MN | V5 | human | -0.26 | 0.42 | 0.001 | <1 |
| 12 | 10 | 2019-05-28 | mAstro | HA | mouse | -0.87 | 0.99 | 0.001 | 158.2 |
| 14 | 10 | 2019-05-28 | MN | V5 | human | -0.31 | 0.23 | 0.017 | 422.8 |
| 20 | 17 | 2019-04-17 | mAstro | HA | mouse | -0.86 | 0.99 | 0.001 | 431.2 |
| 18 | 17 | 2019-04-17 | MN | V5 | human | -0.51 | 0.91 | 0.001 | <1 |
| 28 | 21 | 2019-06-10 | MN | V5 | human | -0.77 | 0.96 | 0.001 | 187.6 |
| 30 | 23 | 2019-06-10 | MN | HA | human | -0.77 | 0.94 | 0.001 | <1 |
| 31 | 25 | 2019-06-17 | MN | V5 | human | -0.84 | 0.96 | 0.001 | <1 |

## Discussion

Our report demonstrates that restricting expression of uniquely-tagged RPL22 (RiboTags) to specific cell types within a mixed co-culture allows for enrichment of cell-type specific translatomes. We fused three commonly used epitope tags (hemagglutinin – HA, V5, and Flag) to the C-terminus of RPL22 and found that V5 and HA, but not Flag RiboTags, facilitated immunoprecipitation of RNA via RNA-bound ribosomes. The failure of RPL22-Flag IP to capture RNA is likely due to inefficient incorporation into the ribosome as indicated by the failure to co-IP RPLP0. Using mixed species co-cultures, we demonstrated that RiboTags can be sequentially IPed without detectable cross contamination by Western blot. Despite efficient separation of RiboTag protein, we observed cross contamination of RNA in RiboTag IPs from mixed species co-cultures. The efficiency of off-target RNA depletion (~80%) was consistent between HA and V5 RiboTags.

We encountered several challenges when applying RiboTags to hiPSCs and hiPSC-derived neural cells. First, constitutive RiboTag expression with the Ef1-alpha promoter was associated with gene silencing in replicative cells (e.g. hiPSCs, NPCs, HEK293s) and reduced viability in post-mitotic cells (e.g. hiPSC-derived neurons). We found that inducible RiboTag expression using the tetracycline response element (TRE) circumvents both of these issues. Approximately twenty-four hours of doxycycline treatment was sufficient for all cell types tested. Reducing the duration or magnitude of induction could minimize potential adverse effects. Promoters with ‘weaker’ activity could also be explored. Second, doxycycline-dependent RiboTag expression is robust in stably transduced hiPSC lines isolated by FACS. However, motor neurons derived from these lines exhibited low and inconsistent expression after the addition of doxycycline. Thus, directed neuronal differentiation may also be associated with gene silencing. RiboTag expression was more robust when neural cells were transduced post dual-SMAD inhibition (see materials and methods). Third, transcription factor overexpression protocols for neural induction frequently use TRE because it is reliable and plasmids are readily available. Although not tested here, we constructed HA and V5 RiboTag vectors flanked by double-floxed inverse orientation (DIO) sequences (Fig. S8) that are compatible with TRE. These vectors could be used in combination with a tamoxifen-inducible Cre or with Cre driven by a cell-type specific promoter^36^.

The challenges outlined above likely contribute to the variability in the depletion efficiency of off-target mRNA. Consistent with differentiation and transduction contributing to technical variability, we observed more consistent reporter gene ratios (mCherry/GFP) across replicates from co-cultures prepared from a single differentiation and transduction per cell type (Fig. S9). However, we cannot rule out that the depletion efficiency may be inherently more variable across hiPSC donors, cell types, or differentiation protocols. Knocking in RiboTags at the native RPL22 locus could reduce variability due to differences in expression and eliminate the need for lentiviral transductions. Such labor-intensive and time-consuming strategies would have to be weighed against the need to include multiple donor hiPSC lines, which is more easily accomplished using lentiviral expression.

RiboTag based enrichment does not eliminate off-target RNA. This may be due to non-specific antibody-RNA interactions or RiboTag-containing ribosomes that bind off-target RNA after cell lysis. Improved purification protocols would need to be compatible across epitope-tags. Whether similar levels of contaminating RNA are observed using RPL10a instead of RPL22 needs to be investigated. Careful consideration should be taken to evaluate the level of contaminating RNA and relative proportion of cells to better inform the downstream analysis. Here we used mixed species experiments to empirically determine the depletion efficiency using off-target RNA. Mitochondrial genes could be used in a similar manner, but would overestimate the depletion efficiency in same species co-cultures because both cell types express the same mitochondrial genes. Here we also used uniquely expressed reporter genes, but these provide only a single datapoint. Computational approaches similar to those used to deconvolve cell-type specific signatures from bulk tissue could be implemented, particularly if information about depletion efficiency is incorporated.

RiboTag and TRAP methods are widely used to investigate cell-type specific transcriptomes *in vivo*. We anticipate that as *in vitro* cultures become increasingly complex in an attempt to more closely mimic *in vivo* cellular networks, methods that disambiguate cell-type specific transcriptomes and avoid the complexity and expense of scRNAseq methods will be necessary. For example, RiboTags could be used to capture cell-type specific RNA from organoids (Fig. S10) and could conceivably be used to capture RNA from multiple cell types simultaneously. scRNA-seq can generate cell-type specific transcriptomes by virtue of profiling thousands of single cells at once. Specific cell types can also be enriched prior to scRNA-seq using FACS-based methods. While the cost and scale of scRNA-seq is rapidly improving, it remains cost-prohibitive for large-scale studies and limited to the most abundant transcripts. RiboTag and TRAP are complementary to scRNA-seq and could even be used in parallel to deeply interrogate full-length transcripts, yielding information about low-abundant transcripts and splicing that are missed by scRNA-seq. Together, these methods are promising avenues for dissecting cell-type specific gene expression in complex *in vitro* cultures.

## Materials and Methods

### Plasmid Construction

All primers and plasmids used in this study are listed in Table S1 and Table S2, respectively, and plasmids will be made available at Addgene. Oligos were purchased from Integrated DNA technologies (IDT). All plasmids were verified by Sanger sequencing. P_Ef1a_-RPL22-3_X_HA_-P2A-_EGFP_-T2A-_Puro was made by Gibson assembly of PCR amplified cPPT/CTS-EF1a (primers JG101/128; addgene #52962 template) and RPL22-3xHA (primers JG106/109; human cDNA template) with EcoRI/PstI digested pLV-TetO-hNGN2_-P2A-_eGFP_-T2A-_Puro (Addgene #79823) backbone. P_Ef1a_-RPL22-3_X_Flag_-P2A-_EGFP_-T2A-_Puro was made by ligating PCR amplified RPL22-3xFlag (primers JG131/132) with EcoRI/AgeI digested P_Ef1a_-RPL22-3_X_HA_-P2A-_EGFP_-T2A-_Puro. P_Ef1a_-RPL22-3_X_HA_-P2A-_EGFP_-T2A-_Neo was generated by Gibson assembly of RPL22-3_x_HA_-P2A-_EGFP (primers JG140/141; RPL22-3_X_HA_-P2A-_EGFP_-T2A-_Puro template) and NotI/XbaI digested pLV-TetO-hNGN2_-P2A-_eGFP_-T2A-_Neo. To make P_Ef1a_-RPL22-3_X_HA_-P2A-_mCherry_-T2A-_Puro, P_Ef1a_-RPL22-3_X_HA_-P2A-_mCherry_-T2A-_Neo, and P_Ef1a_-RPL22-3_X_Flag_-P2A-_mCherry_-T2A-_Puro, mCherry was first amplified from addgene plasmid #22418 (primers JG161/162) and cloned into Topo zero blunt. The resulting plasmid was digested with XbaI and the mCherry fragment was ligated to XbaI digested P_Ef1a_-RPL22-3_X_HA_-P2A-_EGFP_-T2A-_Puro, P_Ef1a_-RPL22-3_X_HA_-P2A-_EGFP_-T2A-_Neo, and P_Ef1a_-RPL22-3_X_Flag_-P2A-_EGFP_-T2A-_Puro, respectively. The 2A peptide sequence causes the ribosome to skip at the 3’ end. Thus, an additional short peptide sequence is added to the end of the epitope tag. To test whether the additional amino acids disrupt the normal function of RPL22, we made P_Ef1a_-EGFP_-T2A-_RPL22-1_x_Flag by Gibson assembly of RPL22_-_1x-Flag (primers JG278/279; RPL22-3_X_HA_-P2A-_EGFP_-T2A-_Puro template), EGFP-T2A-(primers JG276/277; P_Ef1a_-RPL22-3_X_HA_-P2A-_mCherry_-T2A-_Puro template) and EcoRI/AgeI digested addgene #52962. P_Ef1a_-mCherry_-T2A-_RPL22-1_x_V5 was made by Gibson assembly of RPL22-1xV5 (primers JG278/280; RPL22-3_X_HA_-P2A-_EGFP_-T2A-_Puro template), mCherry-T2A (primers 277/281; P_Ef1a_-RPL22-3_X_HA_-P2A-_mCherry_-T2A-_Puro template), and EcoRI/AgeI digested addgene plasmid #52962. To tightly control expression of epitope-tagged RPL22, P_TRE_-mCherry_-T2A-_RPL22-1_x_V5 was generated by Gibson assembly of PCR amplified mCherry-_-T2A-_RPL22-1xV5 (primers JG284/285; template P_Ef1a_-mCherry_-T2A-_RPL22-1xV5) and EcoRI/NheI digested pLV-TetO-hNGN2_-P2A-_eGFP_-T2A-_Puro (Addgene #79823). P_TRE_-EGFP_-T2A-_RPL22-3_x_HA was made by Gibson assembly of GFP-T2A (primers JG290/291; P_Ef1a_-EGFP_-T2A-_RPL22-1_x_Flag template), RPL22-3xHA (JG292/293; RPL22-3_X_HA_-P2A-_EGFP_-T2A-_Puro template), and EcoRI/NheI digested pLV-TetO-hNGN2-P2A-eGFP-T2A-Puro (Addgene #79823). To make Lenti-DIO-P_EF1a-_RPL22-3_x_HA-IRES-YFP, the double-floxed inverse open reading frame (DIO) fragment was amplified from pAAV-Ef1a-DIO-eYFP (gift from Stan McKnight; primers JG303/304) and digested with SpeI. P_Ef1a-_RPL22-3_X_Flag_-P2A-_EGFP_-T2A-_Puro was digested with NheI and treated with Alkaline Phosphatase (AP). The resulting fragments were joined by ligation. Lenti-DIO-P_Ef1a-_mCherry_-T2A-_RPL22-1_x_V5 was made by ligation of PCR amplified mCherry_-T2A-_RPL22-1_x_V5 (primers JG284/285; template P_rTetR-_mCherry_-T2A-_RPL22-1_x_V5) to AscI/NheI digested and AP treated Lenti-DIO-P_EF1a-_RPL22-3_x_HA-IRES-YFP.

### Passaging and maintenance of cell lines

HEK293 and 3T3 cells were maintained in DMEM (Thermo Fisher Scientific) supplemented with 10% Cosmic Calf Serum (Thomas Scientific) and Antibiotic-Antimycotic (Thermo Fisher Scientific). Feeder-free hiPSCs were maintained on Matrigel (Corning) or Geltrex (Thermo Fisher Scientific) coated plates in StemFlex (Thermo Fisher Scientific) as indicated and split either with Accutase (StemCell Technologies) or ReleSR (StemCell Technologies) as recommended by the manufacturer. hiPSC lines used in this study were generated previously^37^ by sendai viral reprogramming^38^. NPCs were cultured in DMEM/F12 supplemented with N2 (Thermo Fisher Scientific), B-27 minus vitamin A (Thermo Fisher Scientific), and FGF-2 (R&D Systems) on Matrigel-coated plates as described^39^.

### Transfections and lentivirus production

Plasmids encoding RiboTag constructs were used to transiently transfect HEK293 cells with lipofectamine 3000 (Thermo Fisher Scientific) as recommended by the manufacturer. Third generation lentiviruses were produced by polyethylenimine transfection with pCMV-VSV-G (addgene #8454), pMDLg/pRRE (addgene #12251), and pRSV-Rev (Addgene #12253), and the appropriate transfer plasmid using established methods ^40^. Lentivirus was concentrated by centrifugation, aliquoted, and stored at −80°C. Prior to use, aliquots of lentivirus were thawed in a water bath at 37°C and added to the appropriate medium.

### FACS

HEK293 cells were dissociated with TrypLE (Thermo Fisher Scientific), washed, resuspended in PBS, and filtered through a 40 μm filter. hiPSCs were dissociated using Accutase (StemCell technologies), washed, resuspended in PBS containing 1% (v/v) BSA, and filtered through a 40 μm filter. Cells positive for mCherry or GFP were isolated by FACS on a SONY SH800. hiPSCs were plated on Geltrex (Thermo Fisher Scientific) or Matrigel (Corning) coated plates in StemFlex supplemented with Thiazovivin (Calbiochem) for 24 hrs.

### Neuron induction and differentiation

hiPSCs were differentiated into MNs according to Maury et al.^41^. A detailed protocol can be found in the supplementary methods and at protocols.io. Briefly, embryoid bodies (EBs) were made by dissociating hiPSCs with Accutase and seeding onto low attachment plates (Corning) in supplemented N2B27. Neural induction occurs via dual-SMAD inhibition^42^ with SB431542 (Stemgent) and LDN-193189 (Stemgent) followed by motor neuron specification by activation of Wnt (Chir-99021; Selleck Chemicals), retinoic acid (RA; Sigma Aldrich), and hedgehog (SAG; Millipore Sigma) signaling. Neuronal maturation occurs in the continued presence of RA and HH cues, but with the addition of DAPT (Tocris), a Notch signaling inhibitor, and GDNF and BDNF to increase neuron survival.

Embryoid bodies were dissociated with 0.25% Trypsin (Thermo Fisher Scientific) in the presence of DNase at 37°C using a thermomixer (Eppendorf) and plated on Geltrex or Matrigel-coated plates. Lentivirus encoding RiboTag constructs was added when plating dissociated EBs or to plated neurons for twenty-four hours.

Induced excitatory neurons were produced by overexpression of NGN2 as described^39^. Briefly, NPCs were seeded onto laminin-coated plates and transduced by spinfection^43^ with *CMV-rtTA* (Addgene ID: 19780) and *TetO-NGN2-P2A-Puro* (Addgene 79049). Where indicated, RiboTag lentivirus was added concurrently. NGN2 and RiboTag were induced with 1μg/ml doxycycline and transduced cells were selected with 1μg/ml puromycin. When needed, media was supplemented with cytosine β-D-arabinofuranoside (Ara-C; Sigma, #C6645) to remove proliferative cells. To avoid RiboTag gene silencing in long-term NGN2-induced neurons, we constructed a lentiviral vector for constitutive expression of NGN2 (P_Ef1a_-NGN2_-T2A-_NeoR) that can be used in combination with tetracycline inducible RiboTag vectors.

Induced GABAergic neurons were produced from NPCs by overexpression of Ascl1 and Dlx2^44,45^. NPCs were transduced by spinfection with lentivirus encoding CMV-rtTA (Addgene 19780), *TetO-Ascl1-T2A-Puro* (Addgene ID: 97329), *TetO-Dlx2-IRES-Hygro* (Addgene ID: 97330) and the indicated RiboTag construct. Doxycycline was added for fourteen days starting 24 hours post-transduction. Transduced cells were selected for five days with puromycin and hygromycin starting 48 hours post transduction. Cells were switched to neuronal media (Neurobasal (ThermoFisher, #21103049) supplemented with Anti-Anti (ThermoFisher, #15240062), N2 (ThermoFisher, 17502-048), B-27 minus vitamin A (ThermoFisher, #12587-010), GlutaMAX (ThermoFisher, #35050061), 1 mg/ml natural mouse laminin (ThermoFisher, #23017-015), 20ng/ml BDNF (Peprotech, #450-02), 20ng/ml GDNF (Peprotech, #450-10), 500 μg/ml cAMP (Sigma, D0627), and 200nM L-ascorbic acid (Sigma, #A0278) on day seven. Half media changes were performed every second day.

### Primary mouse astrocytes

Primary mouse mixed glia cultures were derived from P0 or P1 B6.SJL animals as previously described^46^. Briefly, cortices were dissected and meninges were removed. Tissue was digested in 0.25% Tryspin with EDTA followed by trituration and then strained through a 100 μm strainer. Cell were resuspended in 5 ml of Cortex Glial Media (10% FBS, 1% Pen/Strep, in High Glucose DMEM with Sodium Pyruvate) and plated in T25s coated with 20 μg/ml of Poly-L-Ornithine.

### hiPSC-MN and primary mouse astrocyte co-cultures

RiboTag transduced primary astrocytes were resuspended in motor neuron media (see detailed methods on protocols.io) supplemented with 2% FBS and added to three to four-week old hiPSC-MNs that were previously transduced with a compatible RiboTag construct. hiPSC-MNs and primary mouse astrocytes were co-cultured for at least one week prior to immunoprecipitation.

### Cortical-enriched organoid and microglia co-cultures

Human cortical-enriched organoids (hCO) were made based on the protocol by^47^. Human iPSC lines obtained from the Tau Consortium cell line collection (www.http://neuralsci.org/tau) (GIH7-C2Δ2B12 (MAPT V337V CRISPR corrected to WT/WT), GIH7-C2Δ2A01 (MAPT V337M/WT) and ND32951A.15Δ1B06 (MAPT V337V Crispr corrected to WT/WT), NeuraCell^48^, NY, USA) were maintained in mTeSRTM1 medium (StemCell Technologies, catalog #05851) based on feeder-free culture protocols in six-well plates (Corning, catalog #3506) coated with growth factor-reduced Matrigel (Corning, catalog #356231). At 80-85% confluency, hiPSC colonies were lifted with Accutase (Innovative Cell Tec. #NC9839010), a single cell suspension created, and cells resuspended in E8 medium with ROCK inhibitor Y-27632 (Tocris Cat#1254) at 2 million cells/ml. 3million cells were added per well in an AggreWell™800 plate (Stem Cell Technologies, catalog #34811) (10,000 cells per microwell) and incubated for one day. The resulting spheroids were removed from the microwells and transferred to low-attachment dishes in E6 medium supplemented with 2.5 μM Dorsomorphin (DM) (Tocris, catalog #3093), 10 uM SB431542 (Tocris, catalog #1614), and 2.5 uM XAV-939 (Tocris #3748) to initiate neural differentiation through dual-SMAD inhibition^42^. On day 6, the medium was changed to Neurobasal-A (Life Technologies #10888-022) plus B-27 supplement without vitamin A (Life Technologies, catalog #12587010), GlutaMax (Life Technologies, #3505-061), Antimycotic (Life Technologies, ##15240-062), 20 ng/ml FGF2 (R&D Systems, #233-FB) and 20 ng/ml EGF (Peprotech, # AF-100-15). On day 25, FGF2 and EGF were replaced with 20ng/ml BDNF (Peprotech, # 450-02) plus 20ng/ml NT3 (Peprotech, # 450-03). From day 43 onwards, the medium was changed every four days using only the prior medium without growth factors.At day 20, organoids were QCd by fixing overnight at 4°C in 4% paraformaldehyde (Santa Cruz) followed by sucrose cryoprotection, embedding in OCT and cryosectioning at 20uM thickness, then staining for PAX6 (Biolegend, #901301, 1:200), SOX2 (Santa Cruz, #sc-365823, 1:100), and B-tubulin III (Sigma, # T-8660), 1:1000).

For GIH7-C2-2A01, ND32951A.15-B06, GIH7-C2-2B12 hCO co-cultures, approximately 500,000 human microglia (HMC3, ATCC CRL-3304) transduced with a Ribotag construct were added to 10-month old hCOs in 6-well low attachment plates. hCOs and microglia were co-cultured for 3 weeks prior to immunoprecipitation. 1, 3, or 6 hCOs were pooled for immunoprecipitation. Doxycycline was added 1 day prior to immunoprecipitation.

### Epitope-tagged RPL22 immunoprecipitation

Cycloheximide (CHX) is a translational inhibitor that blocks elongation by inhibiting eEF2-mediated translocation^49^. The tight interaction between the ribosome and mRNA in the presence of CHX allows mRNA to be indirectly purified by immunoprecipitation of epitope-tagged RPL22. A detailed protocol can be found in the supplementary methods. Briefly, cells were washed with PBS and resuspended in ice-cold polysome buffer (50 mM Tris-HCl pH 7.4, 100 mM KCl, 5 mM MgCl_2_, 1 mM DTT) supplemented with CHX (100 μg/ml; Sigma-Aldrich), Turbo DNase (20 U/ml; Thermo Fisher Scientific), Superase-In (400 U/ml; Thermo Fisher Scientific), and protease inhibitor cocktail (Sigma-Aldrich; cat# P8340). Cells were lysed on ice by the addition of NP-40 (1% v/v final concentration) and sodium deoxycholate (0.5% v/v final concentration), cleared by centrifugation, and filtered using a 100 kDa Amicon ultra-2 filter unit (EMD Millipore). Aliquots were removed for input control RNA and Western blot analysis. Anti-HA (Pierce) or anti-V5 (MBL International) magnetic beads were washed with polysome buffer supplemented with Tween (0.1% v/v final concentration), added to the lysate, and incubated for at least 4 hours at 4°C using end over end mixing. Beads were washed with high salt polysome buffer (50 mM Tris-HCl pH 7.4, 300 mM KCl, 5 mM MgCl_2_) supplemented with NP-40 (0.5% v/v final concentration), CHX (100 μg/ml), Superase-In (20 U/ml), and protease inhibitor. For sequential immunoprecipitations, the supernatant was kept and used for an additional immunoprecipitation with the appropriate antibody. For Western blot analysis, beads were resuspended in Laemmli Sample buffer (Bio-rad) supplemented with β-mercaptoethanol (BME) and boiled at 95°C for 5 minutes. For RNA purification, beads were resuspended in Trizol, incubated at room temperature for 3 minutes, and stored at −80°C.

### Western blot

Cell lysates were made as indicated, prepared with Laemmli sample buffer supplemented with BME, and boiled prior to SDS-PAGE. Samples were resolved using 4-20% Mini-PROTEAN TGX precast protein gels (Bio-Rad), transferred to nitrocellulose, blocked with 5% milk in Tris buffered Saline (TBS; 50 mM Tris, 150 mM NaCl) with 0.1% Tween-20 (TBS-T). Primary antibodies include alkaline phosphatase conjugated anti-Flag (Sigma-Aldrich A9469) and anti-HA antibodies (Immunoreagents #MuxOt-111-DALP), biotin conjugated anti-HA (Biolegend 901505) and anti-V5 (Invitrogen MA5-15253-BTIN) antibodies, and unconjugated anti-GFP (Origene TA150070), anti-mCherry (Novus NBP2-25158). Secondary antibodies used include IRDye680 conjugated anti-rabbit (Li-COR 925-68071) and anti-mouse (Li-COR 926-68020) antibodies, and IRDye800 conjugated anti-rabbit (Li-COR 926-32211), anti-mouse (Li-COR 926-32210), anti-chicken (Li-COR 925-32218), and anti-goat (Li-COR 925-32214). Blots probed with alkaline phosphatase conjugated antibodies were visualized using nitroblue tetrazolium (NBT) and 5-bromo-4-chloro-3indolyl phosphate (BCIP) in alkaline phosphatase buffer (100mM Tris-HCl pH 9.5, 5 mM MgCl_2_, 100 mM NaCl). IRDye probed blots were visualized using the Li-COR Odyssey CLx.

### RNA extraction

RNA isolated from lysates prior to immunoprecipitation, referred to as input control RNA, was purified using Trizol LS followed by RNeasy Micro Kit (Qiagen) column purification as follows. 100 μl of lysate was combined with 150 μl of RNase free water and 750 μl of Trizol LS. Immunoprecipitated RNA was eluted from antibody-conjugated beads using Trizol. Samples were then extracted with chloroform, centrifuged, and the aqueous phase from the input control and IP RNA was mixed with 500 μl and 200 μl of 70% ethanol, respectively, and applied to RNeasy MinElute columns (Qiagen). Wash and elution steps were carried out according to manufacturer recommendations. RNA concentration was determined using the Qubit 3.0 fluorometer (Thermo Fisher Scientific). RNA quality was assessed by Bioanalyzer (Agilent).

### Reverse transcription digital PCR (RT-dPCR)

Reverse transcription digital PCR (RT-dPCR) was carried out using a Formulatrix Constellation digital PCR system using a reaction volume of 12 μl in a 96-well format. Applied Biosystems TaqPath One-Step RT-qPCR Master Mix CG was used for all reactions. All reactions were supplemented with 0.1% (v/v) Tween-20. Primers and probes were used at 400nM and 250nM, respectively, for detection of *GFP* and *mCherry.* ACTB primers and probes (IDT reference gene assay – Hs.PT.39a.22214847), were used at 1.5 mM and 750 nM, respectively. Primer and probe sequences are listed in Table S1. Data analysis was performed using R.

### RNA sequencing

RNA was reverse transcribed and amplified using a switching mechanism at the 5’ end of the RNA template ^13^ reaction with Maxima reverse transcriptase (Thermo Fisher Scientific), and oligos listed in Table S1. cDNA yield was measured with a Qubit 3.0 Fluorometer and size distribution was measured by Bioanalyzer (Agilent). Sequencing libraries were constructed using the Nextera DNA Flex kit according to manufacturer instructions. Libraries were quantified by Qubit Fluorometer and qPCR, size distribution measured by Bioanalyzer, and sequenced on an Illumina MiSeq or NovaSeq.

### RNA-seq analysis

RNA-seq reads were processed with TrimGalore (v0.4.4)^50^ and aligned with STAR (v2.5.2a)^35^. For co-cultures composed of human and mouse cells, reads were aligned to a hybrid reference genome that included the canonical chromosomes from hg38 (1-22,X,Y,MT) and mm20 (1-19,X,Y,MT) and sequences for GFP and mCherry. RNA-seq from cultures composed of human cells only were aligned to hg38. At the chromosome level, uniquely mapping reads were quantified with samtools (v1.7). At the gene level, uniquely mapping reads were quantified with featureCounts from the Subread package (v1.5.2)^51^. Count matrices were analyzed in R (v3.6). DESeq2^52^ was used for differential expression analysis Gene set enrichment analysis (GSEA) was carried out using fgsea^53^ using ensembl gene biotype annotation. GSEA for chromosome enrichment was carried out using protein coding genes across the canonical chromosomes.

We calculated the depletion efficiency for each immunoprecipitation as follows. For each library, we counted the number of unique reads mapped to each chromosome and calculated the percentage of the total library represented by each chromosome. We evaluated the relative enrichment for each chromosome compared to the co-culture by plotting the initial percent unique reads (co-culture) against the change in unique reads in the matched IP (IP – coculture). The depletion efficiency for each IP was defined as the slope of the regression line for off-target chromosomes (e.g. mouse chromosomes in a human specific IP and vice versa). The maximum depl

## Oversight

All hiPSC research was conducted under the oversight of the institutional review board (IRB) at the Icahn School of Medicine at Mt. Sinai (ISMMS) and the Embryonic Stem Cell Research Overview (ESCRO) committees at ISMMS and the New York Genome Center (NYGC).

## Data Availability

Donor hiPSC lines used in this study were previously deposited at the Rutgers University Cell and DNA Repository (Study 160; http://nimhstemcells.org). RNA-seq is available at the Gene Expression Omnibus (GEO) repository (accession number: GSE149076) and at https://www.synapse.org/#!Synapse:syn21971188/wiki/602328. Owing to constraints reflecting the original consents, which are restricted to the study of neurodevelopment, the raw RNA-seq data will be made available by the authors upon reasonable request and IRB approval.

## Reagent Availability

All plasmids used in this study will be deposited at Addgene.

## Code Availability

Code is available at https://github.com/jagregory17/RiboTag

## Acknowledgements

This work was supported by Project ALS. ALS sequencing efforts at NYGC are supported by the ALS Association (Grant #19-SI-459) and the Tow Foundation. This work was partially supported by National Institute of Health (NIH) grants R56 MH101454 (K.J.B), R01 MH106056 (K.J.B.) and R01 MH109897 (K.J.B.). We thank Neuracell core facility, NSCI, Rensselaer NY, for generating the organoids and the Tau Consortium and Rainwater Charitable Foundation for their support of organoid production. We thank Thi Vo and Philip De Jager for the HMC3 cell line. We also thank Stan McKnight for the pAAV-Ef1a-DIO-eYFP plasmid.

## Contributions

K.J.B., H.P. and J.A.G. contributed to the experimental design. E.H., R.A., M.H., N.B. and J.A.G. constructed plasmids, lentivirus, and cell lines. E.H., N.B., M.H, G.A., and J.A.G. performed hiPSC cell culture and differentiations. C.B. and M.C. dissected primary mouse astrocytes. E.H., M.H., and J.A.G. performed immunoprecipitations and Western blots. E.H. and J.A.G carried out digital PCR and corresponding analysis. E.H. prepared cDNA and NGS libraries. Organoids were provided by S.T., C.K., A.G., E.C., K.B, S.L., and S.G. J.A.G analyzed NGS data with critical advice from N.S. J.A.G. wrote the manuscript. E.H, G.A. and K.J.B edited the manuscript.

## Ethics Statement

The authors declare no competing interests

## Supplemental Figures

**Figure S1.**
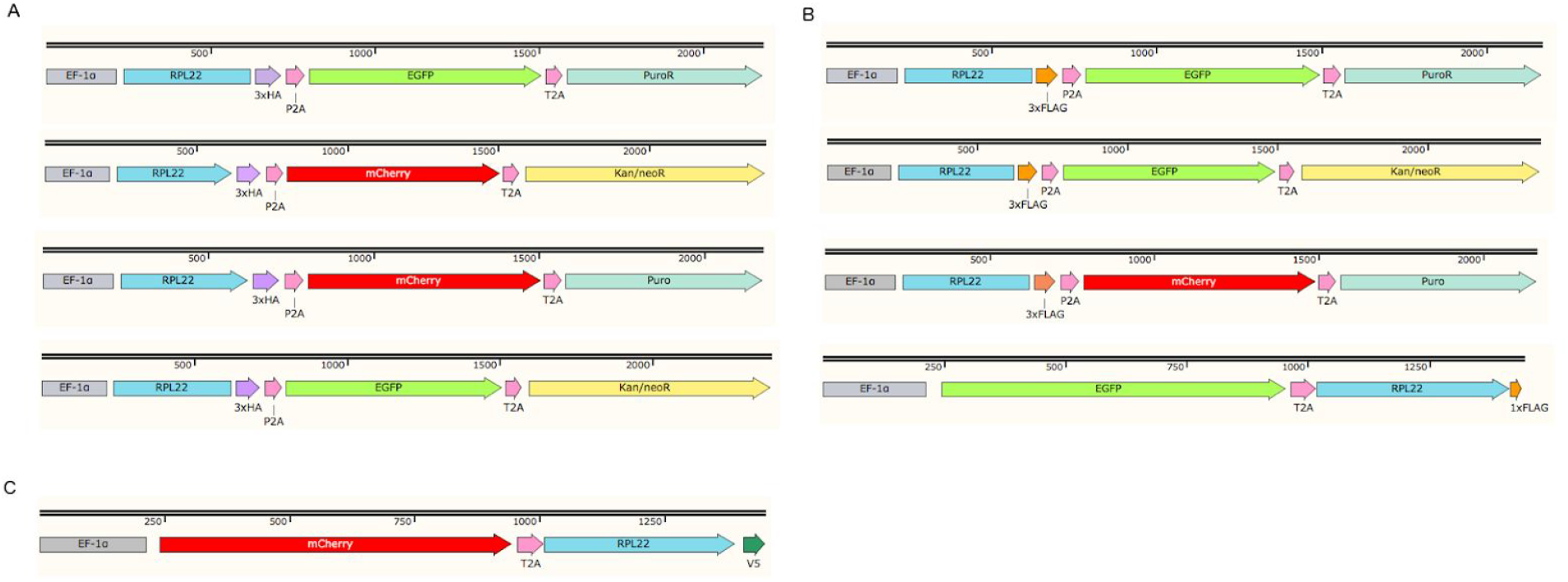
Diagram of RiboTag lentiviral constructs. RPL22 was fused to (A) haemagglutinin – HA, (B) 3xFlag or Flag, or (C) V5 epitope tags. RiboTags are expressed as part of a polycistronic mRNA with a combination of fluorescent reporters and/or selectable markers separated by picornavirus 2A peptides.

**Figure S2.**
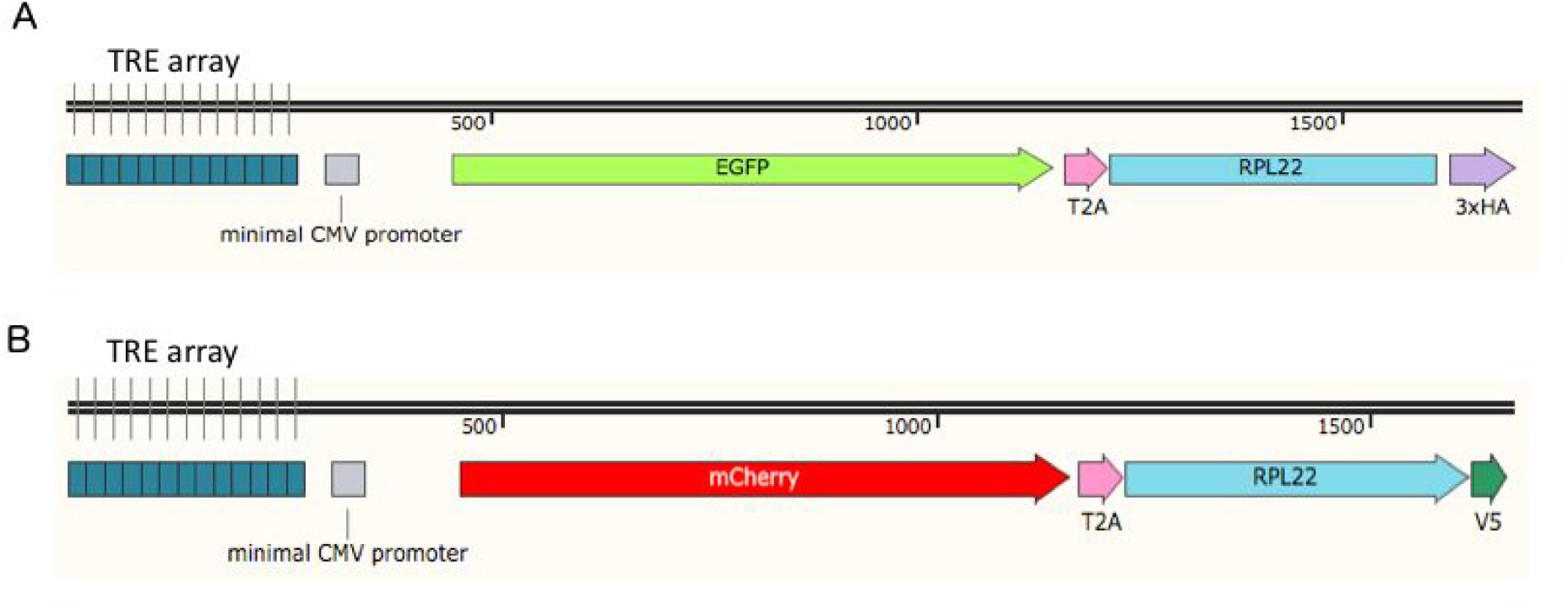
Diagram of inducible RiboTag lentiviral constructs. RPL22 was fused to (A) haemagglutinin – HA or (B) V5 epitope tags. RiboTag expression is induced by the addition of doxycycline. Epitope-tagged RPL22 was moved to the 3’ end of the polycistronic mRNA to minimize extra amino acids in the epitope tag. Tetracycline response element (TRE).

**Figure S3.**
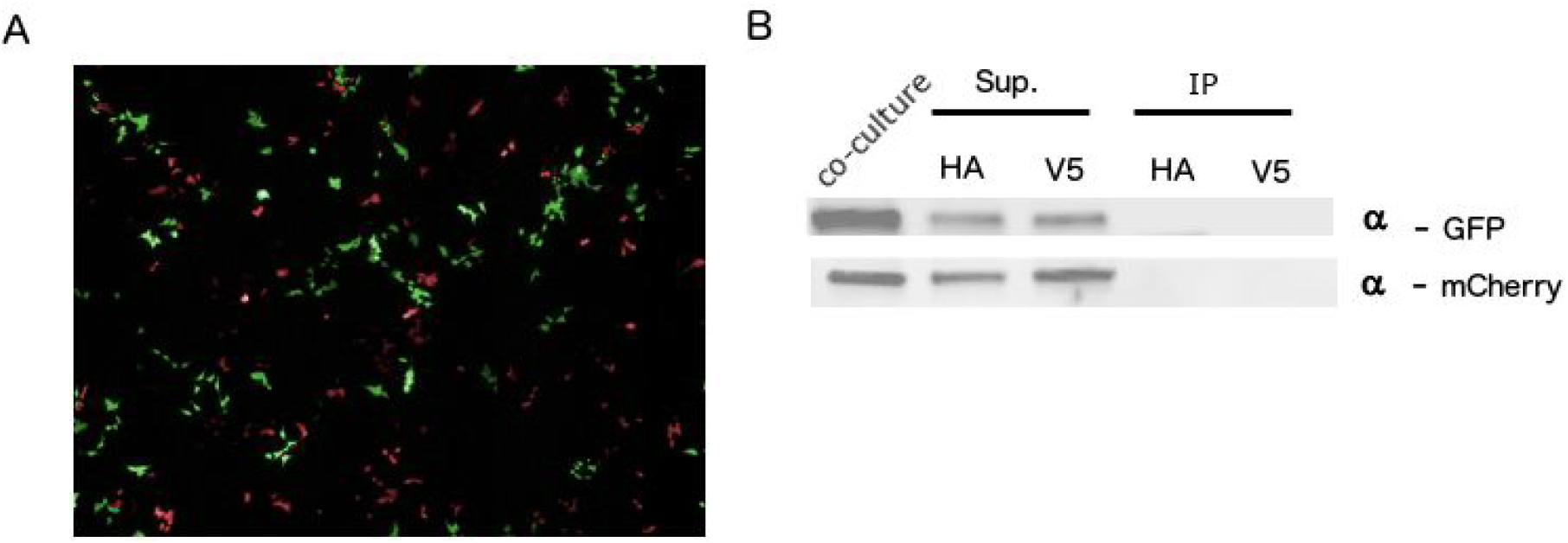
Co-cultures of HEK293 cells transduced with V5 and HA RiboTag vectors. GFP and mCherry protein accumulation in HEK293 cells transduced with PTRE-RiboTag lentivirus were confirmed by (A) microscopy and (B) Western blot. Lysate – total soluble protein. Sup – supernatant. IP – immunoprecipitated protein

**Figure S4.**
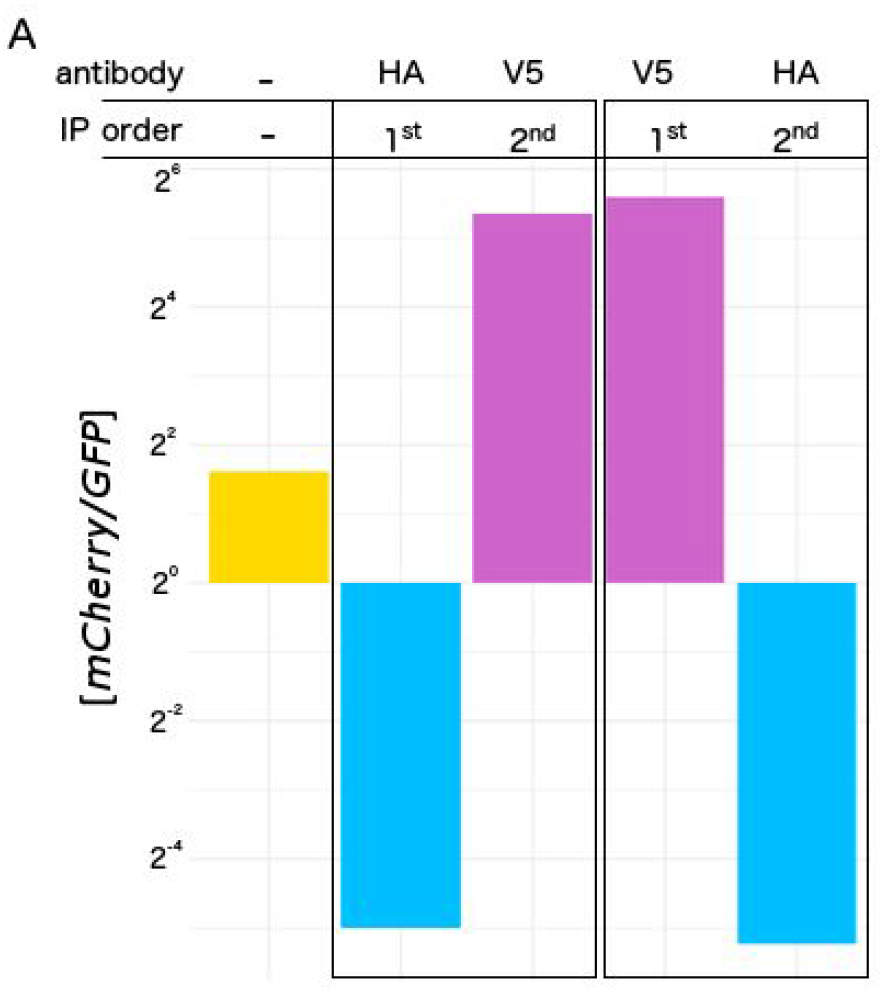
Reverse transcription digital PCR (RT-dPCR) of *mCherry* and *GFP* mRNA concentration from co-cultured human (HEK293) and mouse (NIH-3T3) cells transduced with P_tre_-GFP-T2A-RPL22-HA and P_TRE_-mCherry-T2A-RPL22-V5, respectively. The ratio of *mCherry* to *GFP* transcript abundance was measured in triplicate from pre- and post-IP samples and reported as a ratio of the average concentration. IPs were performed in both orders as indicated.

**Figure S5.**
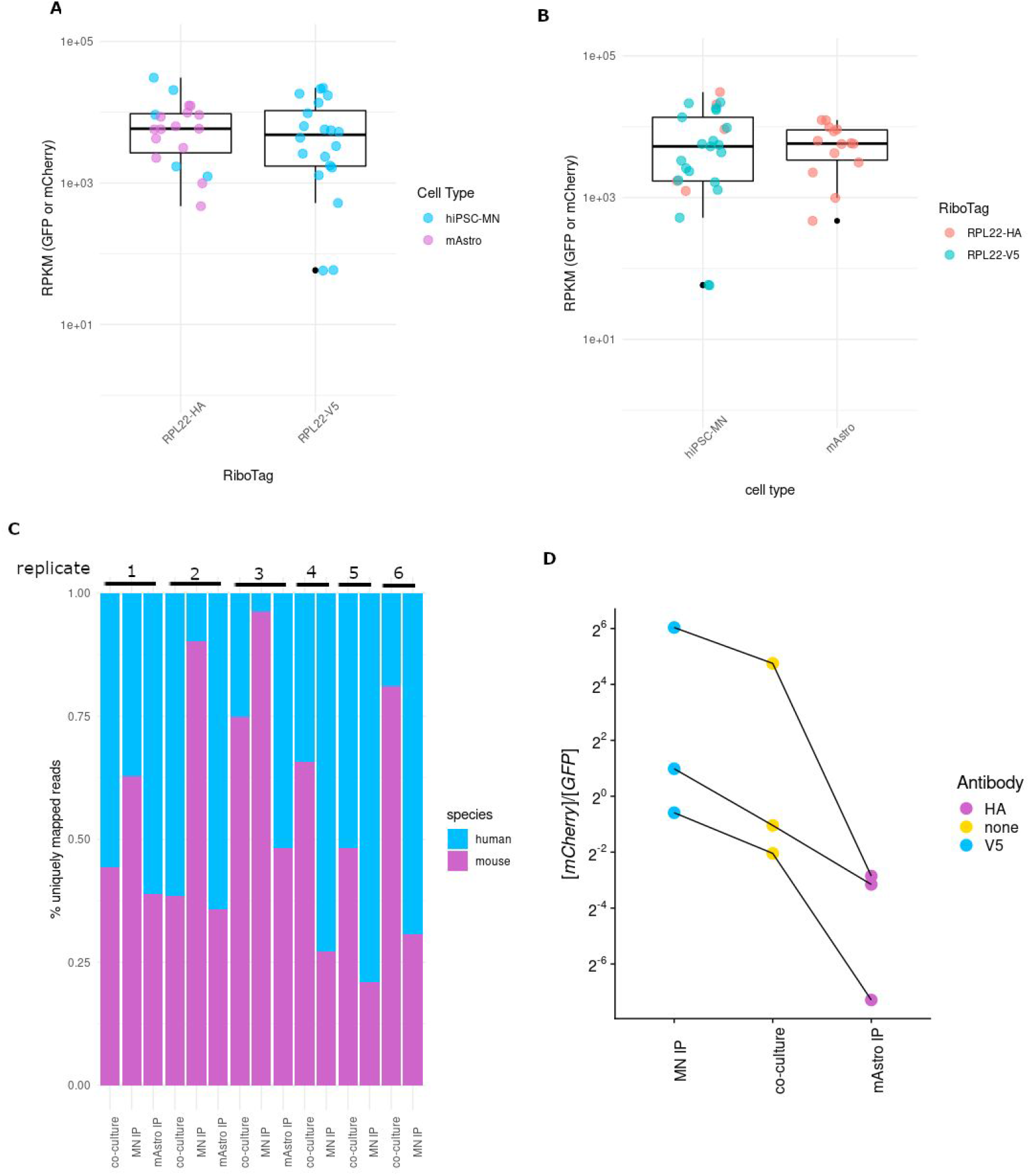
RiboTag expression and cell-type specific enrichment in hiPSC-MN and primary mouse astrocyte co-cultures. (A-B) RPKM of reporter gene expression (GFP/mCherry) in primary mouse astrocytes and hiPSC-derived motor neurons split by (A) RiboTag and (B) cell type. RPKM was calculated using reads mapped to GFP or mCherry and a total library size of reads mapping to the on-target cell type. (C) RNA-seq reads were mapped to a hybrid reference genome containing hg38 and mm20 chromosomes and quantified by species for co-cultures and IP samples. Matched samples are indicated by replicate number. (D) mCherry and GFP levels were measured by RNA-seq. The ratio of mCherry to GFP is reported for co-cultures and IP samples from hiPSC-MNs (MN IP) and primary mouse astrocytes (mAstro IP). HA RiboTag (purple). V5 RiboTag (blue). Lines indicate matched IPs and co-cultures.

**Figure S6.**
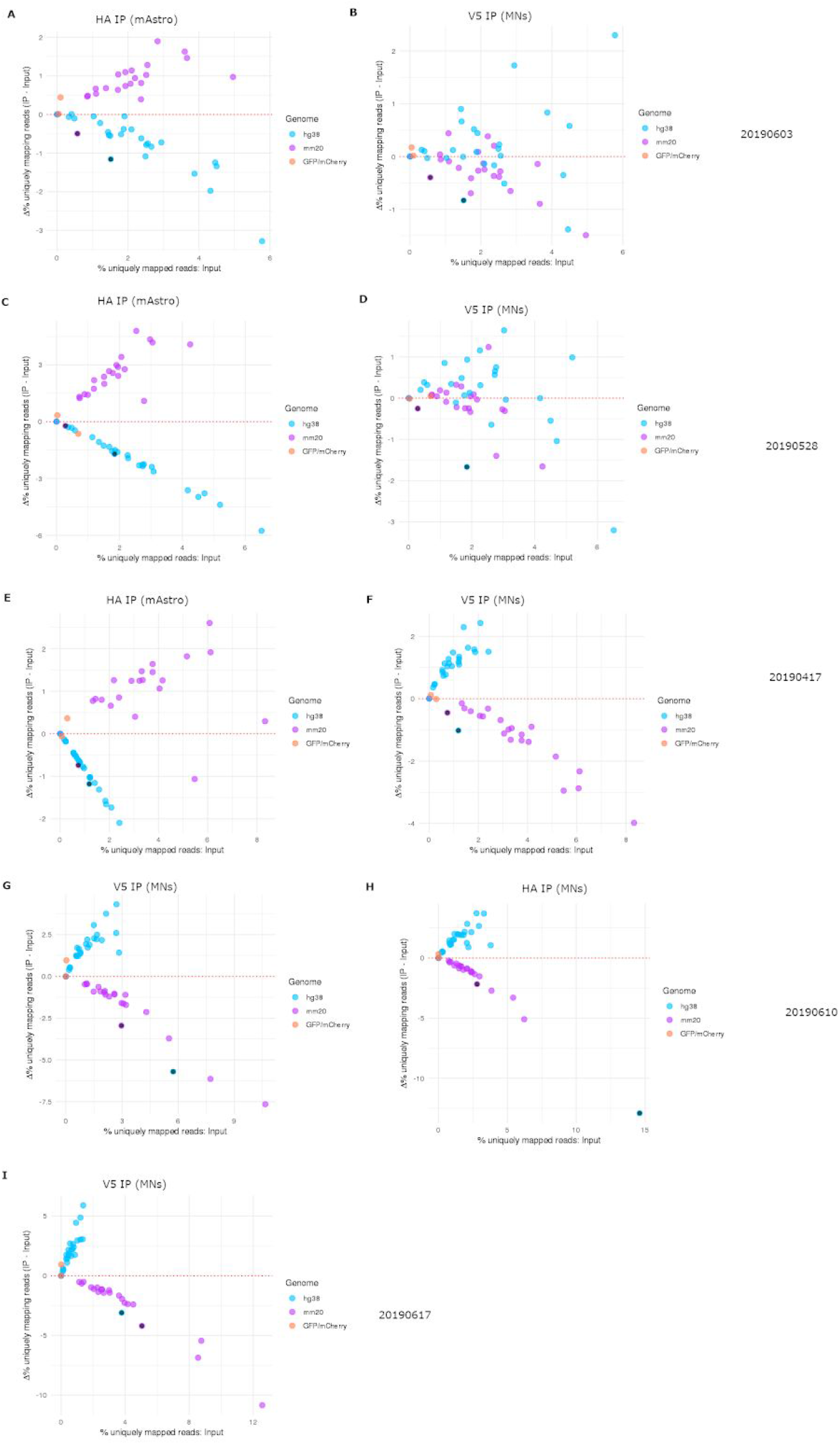
Relative enrichment of human and mouse RNA in RiboTag IPs compared to co-cultures of hiPSC-MNs and primary mouse astrocytes. The change in uniquely mapping reads (IP – input) for each chromosome was plotted against the initial co-culture (Input). Each dot represents a chromosome (hg38 – blue, mm20 – purple; mitochondria are indicated with black fill). Chromosomes that fall above the red dotted line are enriched whereas chromosomes that fall below the red dotted line were depleted in IP samples compared to the Input. (A-B) Matched HA and V5 RiboTag IPs from replicate 1. (C-D) Matched HA and V5 RiboTag IPs from replicate 2. (E-F) Matched HA and V5 RiboTag IPs from replicate 3. (G-I) V5 or HA RiboTag IPs for replicates 4, 5, and 6. Target cell type and date of motor neuron differentiation is indicated for each plot.

**Figure S7.**
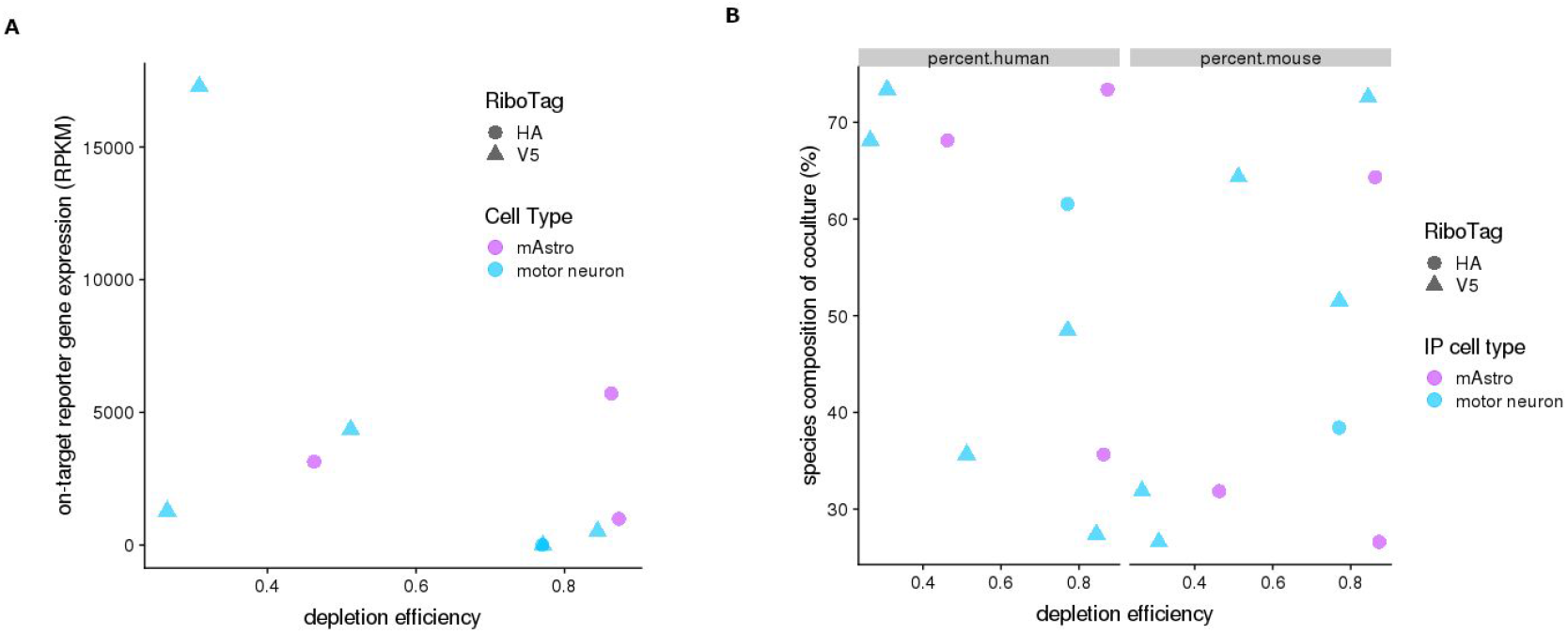
Correlation of RiboTag expression and the initial coculture species composition with depletion efficiency of RiboTag IPs. (A) Scatter plot of on-target reporter (RPL22-HA-GFP; RPL22-V5-mCherry) gene expression in RPKM versus depletion efficiency. RPKM was calculated using the library size for the indicated cell type (i.e. only human reads were used to calculate RPKM for reporter expression in hiPSC-MNs and vice versa). (B) Scatter plot of the initial species composition of the coculture (left – human; right – mouse) versus the corresponding depletion efficiency for every IP. The on-target cell type is indicated by color (mAstro – purple; hiPSC-MN – blue) and shape indicates the RiboTag (HA – circles, V5 – triangles). For example, blue circles & triangles on the left examine the relationship between the depletion efficiency and the relative composition of the on-target cell type (human) in the initial coculture. The purple circles on the left examine the relationship between the depletion efficiency and the off-target cell type (mouse). The reverse is true for the plot on the right.

**Figure S8.**
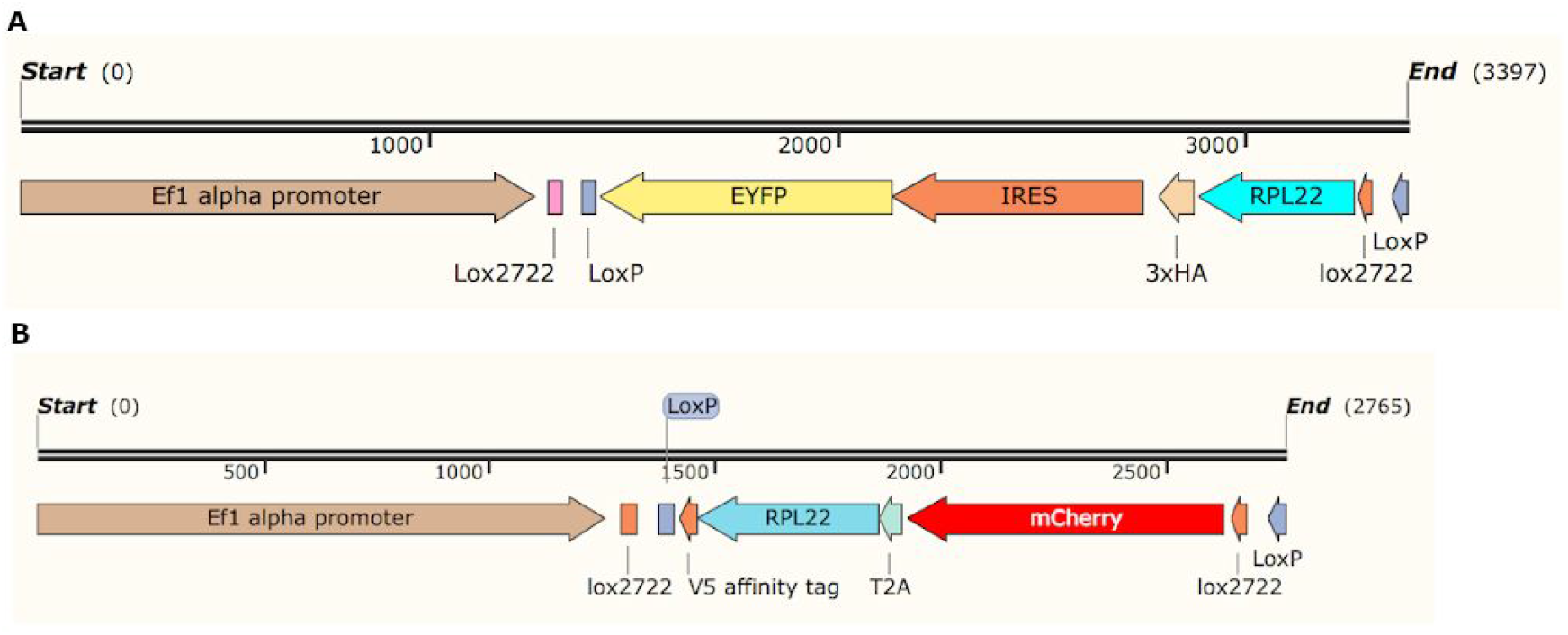
Diagram of double-floxed inverse orientation (DIO) lentiviral RiboTag constructs. (A) EFIa-DIO RPL22-IRES-YFP was amplified from pAAV-Ef1a-DIO-Rpl22-3xHA-IRES-eYFP (gift from Stan McKnight) and cloned into existing RiboTag lentiviral vectors (B) DIO mCherry-T2A-RPL22-V5 RiboTag. RiboTag expression is induced in the presence of Cre, which flips the RiboTag orientation to match the Ef1a promoter.

**Figure S9.**
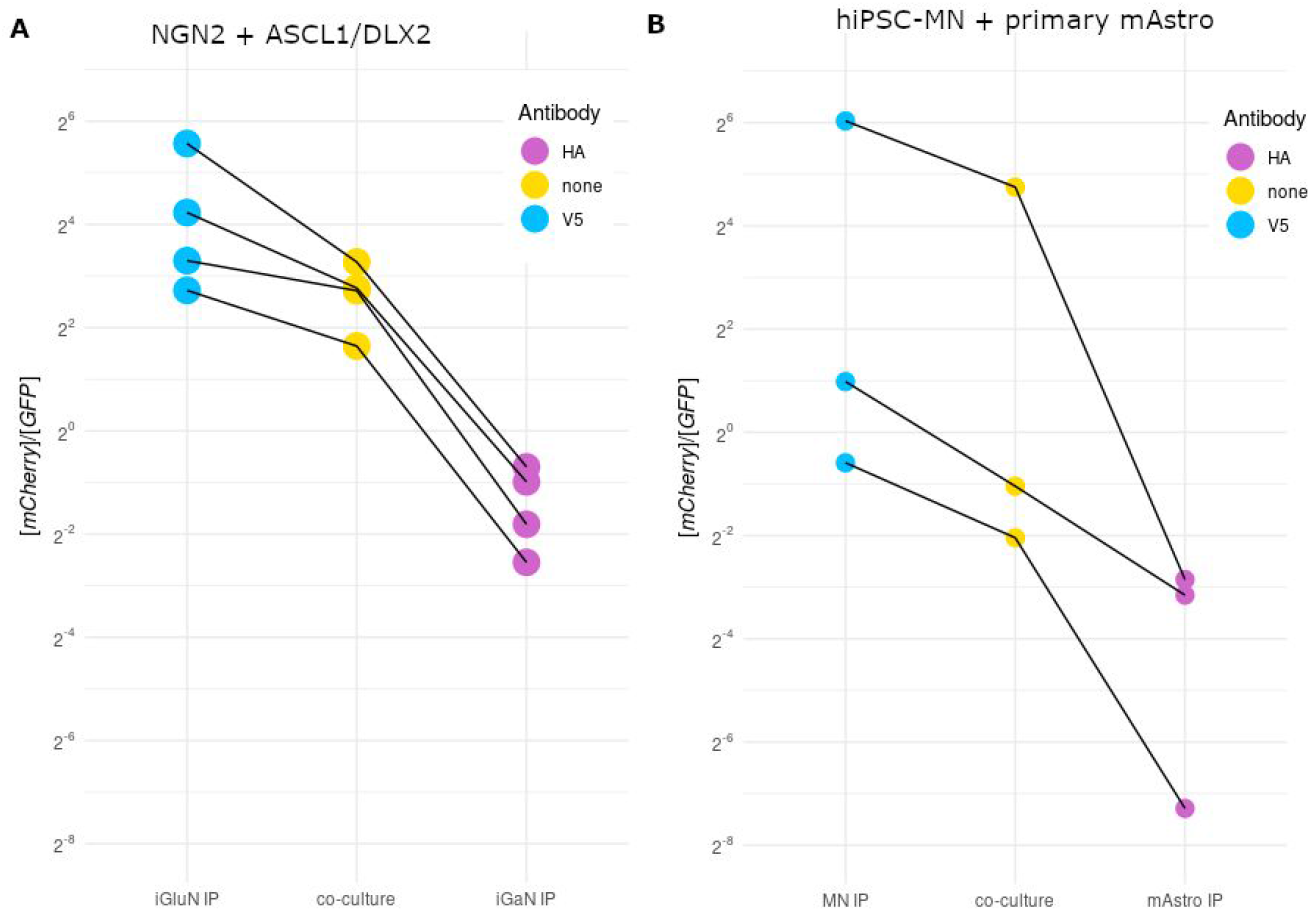
Reverse transcription digital PCR (RT-dPCR) of mCherry and GFP from RiboTag co-cultures. Plotted is the ratio of mCherry to GFP across co-cultures (yellow), V5 IPs (blue), and HA IPs (purple) for (A) NGN2-induced excitatory neuron (iGluN) and ASCL1/DLX2-induced GABAergic neuron (iGaNs) co-cultures prepared from a single differentiation and RiboTag transduction per cell type, and (B) primary mouse astrocytes (mAstro) and hiPSC-MN co-cultures across independent replicates. Lines indicated matched IPs and co-cultures.

**Figure S10.**
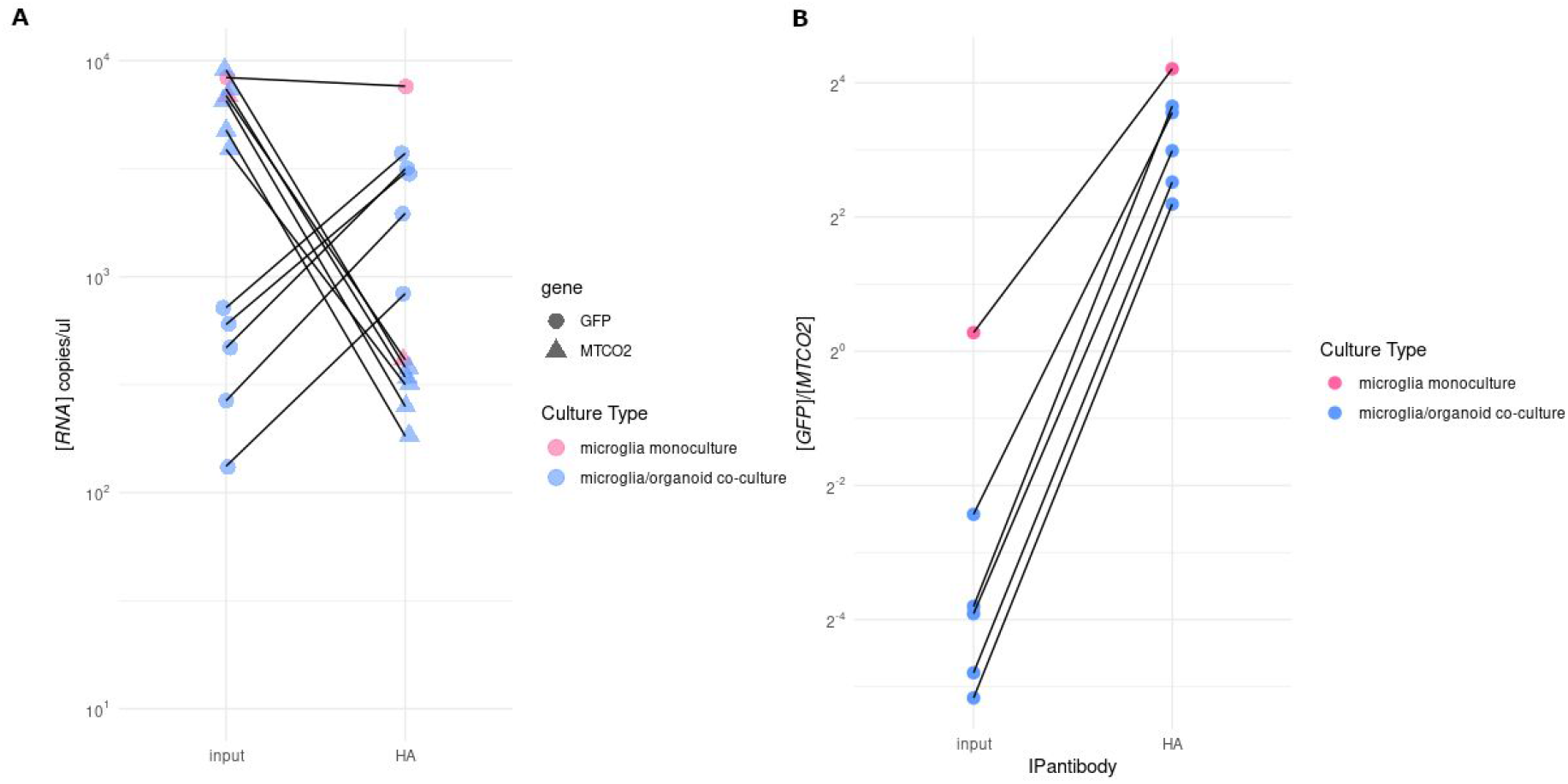
Reverse transcription digital PCR (RT-dPCR) of GFP and MTCO2 from HMC3 microglia expressing GFP-T2A-RPL22-HA co-cultured with neurospheres. Input and HA-immunoprecipitated RNA was purified from stable HMC3 microglia grown alone and in co-culture with neural organoids. GFP and MTCO2 were measured by RT-dPCR and (A) plotted as absolute RNA levels and (B) a ratio of GFP to MTCO2. Lines indicate matched input and IPed samples.

